# Nitrogen and oxygen isotope effects during enzymatic nitrate reduction *in vitro* and by natural lake water consortia

**DOI:** 10.1101/2025.03.29.645582

**Authors:** Jana Tischer, Jakob Zopfi, Claudia Frey, Jessica Venetz, Elena Giglio, Lukas Burgdorfer, Oliver Rehmann, Scott D. Wankel, Moritz F. Lehmann

## Abstract

Nitrate (NO_3_^-^) isotope ratios are useful indicators for nitrogen (N)-transformation processes if the associated isotope effects and their environmental controls are well-constrained. The NO_3_^-^ isotope effects in natural environments may depend on the type of dissimilatory nitrate reductases involved and the degree of isotopic overprinting via NO_3_^-^ regeneration. We measured the coupled N and oxygen (O) isotope effects of NO_3_^-^ reduction in anoxic incubation experiments with laboratory cultures (*Pseudomonas* sp. and *Escherichia coli*) harboring different nitrate reductase enzymes (Nar and/or Nap) as well as with natural freshwater consortia from Lake Lugano (Switzerland) and Lake La Cruz (Spain). For comparison, isotope effects were also evaluated through coupled N and O isotope measurements in the redox transition zone of Lake Lugano North Basin. Incubation-based Rayleigh model N isotope effects (ε_N_) were variable, ranging from 9 to 30‰. In comparison, in situ ε_N_ values (5 to 14‰) estimated by the closed system model were lower, likely due to substrate limitation in the water column. Experiments with *Pseudomonas* sp. and *E. coli* cultures possessing Nar yielded N isotope effects of similar magnitudes and, consistent with previous data, robust Δδ^18^O:Δδ^15^N enrichment ratios of ∼0.9 - 1.0. Nitrate reduction by cultures possessing solely Nap led to lower Δδ^18^O:Δδ^15^N of ∼0.7. In anoxic incubations of lake water, where the effect of nitrification could be excluded, Δδ^18^O:Δδ^15^N values between 0.6 and 1.0 suggest "community activity" of both Nar and Nap. Interestingly, stimulation of lithotrophic nitrate reduction in incubations with amended NO_3_^-^ + sulfide resulted in a Δδ^18^O:Δδ^15^N slope of 0.90 ± 0.03 (standard error, SE), indicating Nar as the more dominant nitrate-reducing enzyme. On the contrary, stimulation of organotrophic nitrate reduction in NO_3_^-^ + acetate amended incubations resulted in a significantly lower slope of 0.72 ± 0.03 SE, suggesting a greater contribution by Nap. In contrast to the nitrate-reduction incubation experiments, the in situ Δδ^18^O:Δδ^15^N value of 1.36 ± 0.14 SE observed in the Lake Lugano water column was unusually high for a freshwater environment, likely reflecting the superimposed effect of NO_3_^-^ production by nitrification. Our study thus underscores that both variations in activity of Nar versus Nap during nitrate reduction, as well as isotopic overprinting by nitrate regeneration may impact ecosystem NO_3_^-^ isotope dynamics in natural denitrifying environments.

## 1. Introduction

The large-scale application of nitrogen (N)-based artificial fertilizers in agriculture, has introduced vast amounts of fixed N into natural habitats, including freshwater and coastal marine environments, where it can cause eutrophication (Gruber and Galloway, 2008). In this context, along the freshwater-marine continuum, lakes fill a crucial role in removing fixed N species, such as nitrate (NO_3_^-^), before they reach the ocean, with denitrification as the most important fixed N sink (Harrison et al., 2009). During this process, microorganisms reduce NO_3_^-^ to nitrite (NO_2_^-^), nitric oxide (NO), nitrous oxide (N_2_O) and ultimately dinitrogen gas (N_2_), under anoxic to low-oxic (O_2_) conditions, allowing its release into the atmosphere. Other N-transformation processes, such as the dissimilatory nitrate reduction to ammonium (DNRA) and nitrification, the oxidation of ammonium (NH_4_^+^) to NO_2_^-^ and NO_3_^-^, retain fixed N in the system (Schlesinger and Bernhardt, 2012). Nitrate reduction as well as other N-cycle reactions exhibit significant N (and O) isotope fractionation (Lehmann et al., 2003; Granger et al., 2008; Wenk et al., 2014). Hence, the isotopic composition of dissolved N species, especially of NO_3_^-^, can be a valuable tool to identify N-transformation processes in natural environments, if the isotope effects (both at the enzyme and ecosystem level) specific to individual N cycling pathways are well constrained.

Isotopic fractionation during unidirectional biological reactions occurs when enzymes transform different isotopocules of a substrate at different rates. Usually, in enzymatic reactions, the lighter molecules are preferentially transformed (i.e., at faster rates than for the heavier molecules), leaving the remaining substrate pool enriched, and the product depleted, in the heavy isotopes (i.e., normal fractionation). Reactions with the faster transformation of heavier molecules result in inverse isotope effects. Isotope ratios are reported in the conventional delta notation as per mil (‰) deviation from the isotopic composition of atmospheric air (AIR) for N, and standard mean ocean water (VSMOW) for O, with δ^15^N = ([^15^N/^14^N]_sample_/[^15^N/^14^N]_AIR_ - 1) x 1000 and δ^18^O = ([^18^O/^16^O]_sample_/[^18^O/^16^O]_VSMOW_ - 1) x 1000. The kinetic isotope effect, ε, is a measure of the strength of isotope discrimination, also expressed in per mil. It is defined as ([*k*_light_/*k*_heavy_] - 1) x 1000, where *k*_light_ and *k*_heavy_ are the reaction rate coefficients for the light and heavy isotopes, respectively. A wide range of ^15^ε and ^18^ε values for nitrate reduction to nitrite, between 3 and 40‰ and between 2 and 33‰, respectively, has been reported in numerous field and culture studies with denitrifying bacterial isolates (Mengis et al., 1999; Granger et al., 2008; Asamoto et al., 2021; summary in Table S1). The high variability in N and O isotope effects may arise from changes in the ratio of enzymatic NO_3_^-^ reduction to cellular NO_3_^-^ transport and may be influenced by factors such as nitrite accumulation, cell-specific nitrate reduction rates, and/or carbon source (Granger et al., 2008; Kritee et al., 2012; Frey et al., 2014b). In natural aquatic environments near redox transition zones (RTZs), the stable isotopic composition of the nitrate and nitrite (i.e. NO_x_) pools may also be affected by oxidative processes (i.e., nitrification), which fractionate stable isotopes as well. The N isotope effect of NH_4_^+^ oxidation to NO_2_^-^, for example, ranges between ∼12 and 38‰ (Denk et al., 2017). In contrast, inverse N isotope effects have been reported for NO_2_^-^ oxidation to NO_3_^-^ by nitrifying (-13‰) or anaerobic ammonium oxidizing (anammox; -31‰) bacteria (Casciotti 2009).

Studies on coupled nitrate N and O isotope shifts during dissimilatory denitrification have revealed that net nitrate reduction does not always impart proportional heavy N and O isotope enrichments in the nitrate pool, and that robust deviations from full proportionality may reflect the type of denitrification and/or the involvement of other biogeochemical N cycling reactions (Granger et al., 2008; Granger and Wankel, 2016). Indeed, the nitrate O versus N isotope enrichment ratio, expressed as Δδ^18^O:Δδ^15^N (≈ ^18^ε:^15^ε) (Granger et al., 2008), seems to depend on two major factors, i) the active nitrate reductase involved and ii) possible isotope overprinting via nitrite oxidation during nitrification or anammox (Granger et al., 2008; Granger and Wankel, 2016). Two dissimilatory nitrate reductases, the membrane-bound Nar, encoded by the *narG* gene, and the periplasmatic Nap, encoded by *napA*, are known enzymes (Sparacino-Watkins et al., 2014) that are used by organotrophic organisms and have been associated with either denitrification or DNRA (Granger et al., 2008; Frey et al., 2014b; Pandey et al., 2020; Asamoto et al., 2021). Pure culture studies with organotrophic denitrifying organisms using only Nar produce robust Δδ^18^O:Δδ^15^N slopes of ∼0.9 to 1, while denitrification by Nap appears to produce systematic Δδ^18^O:Δδ^15^N ratios of ∼0.5 to 0.7 (Granger et al., 2008; Asamoto et al., 2021). Isotopic overprinting of the NO_3_^-^ pool by nitrate production depends primarily on the N and O isotopic composition of NO_2_^-^. The NO_2_^-^ pool is affected by the production mechanism (i.e., nitrification or nitrate reduction), and hence by the isotope ratios of the precursor molecules, by fractional NO_2_^-^ consumption (both by reductive and/or oxidative processes), and, in the case of O isotopes, by O-atom exchange between NO_2_^-^ and water (H_2_O) (Kumar et al., 1983; Buchwald et al., 2012). Depending on the ratio of nitrate production versus reduction rates, and depending on the δ^18^O of the ambient H_2_O, which is significantly lower in freshwater than in marine water, nitrification can either lower or increase the Δδ^18^O:Δδ^15^N beyond slopes of 1:1, as shown in a numerical modelling study (Granger and Wankel, 2016). In marine environments, where reported Δδ^18^O:Δδ^15^N ratios are always ≥1 (Table S1), they are assumed to represent either a pure Nar signal (∼1) or a Nar signal with isotopic overprinting by nitrification (>1) (Sigman et al., 2005; Lao et al., 2019). However, in freshwater, where reported Δδ^18^O:Δδ^15^N ratios mostly range between 0.5 and 1 (e.g., Lehmann et al., 2003 and references therein), it remains a conundrum, as to whether the low values are the result of a higher relative importance of Nap, or of isotopic overprinting, or both (Wenk et al., 2014; Granger and Wankel, 2016).

Lake Lugano is an anthropogenically impacted lake with a history of high N inputs and eutrophication. The North Basin of the lake can be characterized as meromictic (Holzner et al., 2009; Lehmann et al., 2015). Similarly, Lake La Cruz is a meromictic lake, yet without anthropogenic eutrophication. Previous work has revealed that the RTZ of both lake basins acts as an efficient sink of fixed N, and as a hotspot of anaerobic and micro-aerobic N-transforming processes (i.e., organotrophic and S-dependent denitrification, DNRA, anammox, and nitrification in Lake Lugano; denitrification and nitrification in Lake La Cruz) occurring in close proximity (Wenk et al., 2013; Tischer et al., 2022). Hence, both lake basins represent excellent model systems, in which to investigate the interactive nitrate N and O isotope effects imparted by nitrate reduction and production.

Here, we investigated the controls on N and O nitrate isotope effects in net-denitrifying freshwater settings, focusing on the potential of Δδ^18^O:Δδ^15^N as an indicator of enzymatic nitrate reduction versus nitrate regeneration. We used a multifaceted approach: (1) Culture experiments with *Pseudomonas* sp. and *Escherichia coli*, harboring *narG* and/or *napA*, were conducted to extend our knowledge of enzyme-specific nitrate N and O isotope effects associated with nitrate reduction. (2) Anoxic incubation experiments with natural N-reducing microbial communities from the water columns of Lake Lugano and Lake La Cruz were performed to gain insight into the community nitrate isotope effects of nitrate reduction, but under conditions that should eliminate any isotopic overprinting by nitrification. We specifically aimed to evaluate whether observed nitrate N and O isotope effects depend on the metabolism of the active microorganisms (i.e., organotrophic versus lithotrophic N-reduction, possibly involving the differential use of Nar versus Nap, respectively). (3) Finally, we wanted to constrain the impact of Nar-versus-Nap activity and isotopic overprinting by nitrate regeneration on in situ dual O-versus-N nitrate isotope signatures in a natural water column. Here we concentrated on nitrate isotope measurements across the RTZ of the Lake Lugano North Basin.

## 2. Methods

### 2.1. Pure culture incubation experiments

We determined the N and O isotope effects of nitrate reduction in cultures of different strains of *Pseudomonas* sp. and *Escherichia coli* harboring *narG* and/or *napA* (Table 1). *Pseudomonas* sp. strains were isolated from the rhizosphere of perennial grasses (Roussel-Delif et al., 2005). Strain DLT1NRS3 harbors only *narG* and ELC1RS10 both *narG* and *napA*. Strain ELC2NRS8 was reported to possess *napA* only (Roussel-Delif et al., 2005), but we also detected *narG* using the same reported protocol and primers of (Roussel-Delif et al., 2005). In a first set of experiments (“Exp_culture_P1”), the three *Pseudomonas* sp. strains were preincubated in Tedlar Bags (Granger et al., 2008) with modified Burke medium (Atlas, 2010) with yeast extract and tryptone/peptone from casein and without NO_3_^-^. Oxygen in the preincubations was consumed completely overnight, which was confirmed as the bacteria immediately started consuming NO_3_^-^, once we started the incubation by adding 0.5 or 1 mM KNO_3_ (final concentration) from a sterile, anoxic 10 mM stock solution. In the second set of experiments (“Exp_culture_P2”), we incubated *Pseudomonas* sp. strain ELC2NRS8 with glucose, sodium citrate, or sodium succinate, after identifying these substrates as suitable carbon source via GN2-Microplates (Biolog Inc.). Here, the bacteria were incubated in 1-L borosilicate bottles for up to 3 days at 30 °C in Burke medium with the respective carbon source (∼2.5 mM final concentration), after 200-300 µM NO_3_^-^ (final concentration) was added from a 41 mM sterile stock solution. The medium had been purged with N_2_ for 30 minutes to create anoxic conditions. In a third set of experiments (“Exp_culture_Ecoli”), we investigated nitrate reduction by *E. coli* strains JCB4011 (*narG)* and JCB4031 (*napA*) (Potter et al., 1999). The cultures were pre-incubated in lysogeny-broth medium and subsequently incubated for up to 30 h at 37 °C and 70 rpm in 1-L borosilicate bottles containing Burke medium, glycerol, and initial NO_3_^-^ concentrations between 73 and 292 µM. For all incubations, we collected subsamples for NO_3_^-^ and NO_2_^-^ concentration and NO_3_^-^ isotope analysis at regular time intervals through 0.45 µm syringe filters, respectively. Isotope samples were treated with sulfamic acid to remove nitrite (Granger and Sigman, 2009). Samples for concentration analysis were stored at 4 °C, whereas samples for isotope analyses were stored frozen (-20 °C).

**Table 1.** Nitrate isotope fractionation for nitrogen (^15^ε) and oxygen (^18^ε), and Δδ^18^O:Δδ^15^N ratios during pure culture experiments with *Pseudomonas* sp. and *Escherichia coli* strains harboring *narG* and/or *napA*, respectively. Data are presented with standard error of means. Non-significant values are indicated by “ns”.

| Organism | Strain | Carbon source | Nitrate reductase gene | [NO<sub>3</sub><sup>-</sup>]<br>initial (μM) | $^{15}\epsilon$<br>(‰) | $^{18}\epsilon$<br>(‰) | $\Delta\delta^{18}\text{O}:\Delta\delta^{15}\text{N}$ | NO<sub>3</sub><sup>-</sup> reduction<br>(μmol L<sup>-1</sup> h<sup>-1</sup>) | NO<sub>2</sub><sup>-</sup> production<br>(μmol L<sup>-1</sup> h<sup>-1</sup>) |
| --- | --- | --- | --- | --- | --- | --- | --- | --- | --- |
| *Pseudomonas* sp. | DLT1NRS3 | Yeast extract,<br>tryptone/peptone | *narG* | 459 | 12.5 ± 2.2 | 12.3 ± 1.5 | 0.94 ± 0.05 | 29.5 ± 6.1 | 17.5 ± 2.7 |
| *Pseudomonas* sp. | DLT1NRS3 | Yeast extract,<br>tryptone/peptone | *narG* | 511 | 21.9 ± 1.7 | 20.6 ± 1.4 | 0.95 ± 0.02 | 78.9 ± 21.4 | 68.0 ± 16.6 |
| *Pseudomonas* sp. | ELC2NRS8 | Yeast extract,<br>tryptone/peptone | *narG* & *napA* | 823 | 17.4 ± 0.4 | 17.1 ± 0.7 | 0.98 ± 0.02 | 144.5 ± 48.3 (ns) | 160.2 ± 66.9 (ns) |
| *Pseudomonas* sp. | ELC2NRS8 | Yeast extract,<br>tryptone/peptone | *narG* & *napA* | 582 | 22.4 ± 0.5 | 21.0 ± 0.4 | 0.94 ± 0.02 | 75.3 ± 11.5 | 52.5 ± 11.0 |
| *Pseudomonas* sp. | ELC1RS10 | Yeast extract,<br>tryptone/peptone | *narG* & *napA* | 823 | 22.9 ± 0.7 | 21.5 ± 0.9 | 0.94 ± 0.02 | 157.7 ± 34.7 (ns) | 187.0 ± 52.8 (ns) |
| *Pseudomonas* sp. | ELC1RS10 | Yeast extract,<br>tryptone/peptone | *narG* & *napA* | 681 | 21.3 ± 2.6 | 20.6 ± 2.6 | 0.97 ± 0.01 | 102.1 ± 20.7 | 60.1 ± 9.3 |
| *Pseudomonas* sp. | ELC1RS10 | Yeast extract,<br>tryptone/peptone | *narG* & *napA* | 544 | 22 ± 0.5 | 21.3 ± 0.4 | 0.96 ± 0.01 | 61.5 ± 8.7 | 51.8 ± 7.3 |
| *Pseudomonas* sp. | ELC2NRS8 | Citrate | *narG* & *napA* | 227 | 7.9 ± 1.5 | 8.5 ± 1.2 | 0.99 ± 0.17 | 3.0 ± 0.4 | 1.4 ± 0.3 |
| *Pseudomonas* sp. | ELC2NRS8 | Citrate | *narG* & *napA* | 252 | 25.0 ± 3.0 | 22.5 ± 3.2 | 0.90 ± 0.02 | 1.9 ± 0.5 | 0.0 ± 0.0 |
| *Pseudomonas* sp. | ELC2NRS8 | Glucose | *narG* & *napA* | 268 | 30.2 ± 0.9 | 28.2 ± 0.7 | 0.93 ± 0.01 | 1.9 ± 0.6 | 0.6 ± 0.1 |
| *Pseudomonas* sp. | ELC2NRS8 | Succinate | *narG* & *napA* | 262 | 27.0 ± 1.4 | 25.0 ± 1.4 | 0.92 ± 0.01 | 2.0 ± 0.6 | 0.0 ± 0.0 |
| *E. coli* | JCB4011 | Glycerol | *narG* | 249 | 14.9 ± 0.6 | 14.7 ± 0.3 | 0.98 ± 0.12 | 31.1 ± 0.8 | 26.7 ± 1.1 |
| *E. coli* | JCB4011 | Glycerol | *narG* | 81 | 16.7 ± 1.3 | 16.4 ± 1.6 | 0.98 ± 0.02 | 11.7 ± 2.5 | 16.1 ± 2.0 |
| *E. coli* | JCB4031 | Glycerol | *napA* | 292 | 9.6 ± 0.7 | 6.4 ± 0.3 | 0.65 ± 0.02 | 4.6 ± 0.5 | 3.5 ± 0.3 |
| *E. coli* | JCB4031 | Glycerol | *napA* | 73 | 8.0 ± 0.4 | 5.9 ± 0.3 | 0.74 ± 0.03 | 4.7 ± 0.7 | 2.7 ± 0.4 |

### 2.2. Study sites and sampling

*Lake Lugano* - Lake Lugano is a sub-alpine lake on the Swiss-Italian border, situated at an altitude of 271 m above sea level. A natural dam divides the lake into the seasonally mixing South Basin and the semi-permanently stratified North Basin, which has a maximum depth of 288 m. Over the past 50 years, mixing events only occurred in 2005 and 2006, when cold and windy winters caused the complete overturn of the water column (Holzner et al., 2009; Lehmann et al., 2015). In this study, samples were collected in the North Basin at the deepest point (46°00’37.7"N, 9°01’14.9"E). Using 5-L Niskin bottles, we took discrete water samples for nutrient and nitrate isotope analysis in the RTZ between 75 m and 165 m during 12 sampling campaigns in 2015 (February, April, June, August, October, December), 2016 (March, September, November), 2017 (February, October), and 2018 (April). We used a conductivity, temperature, depth (CTD) probe (Idronaut Ocean Seven 316Plus) with an oxygen sensor to determine O_2_ concentrations. For hydrogen sulfide (H_2_S) analyses, we fixed unfiltered water samples with zinc acetate (ZnAc) and monobromobimane, respectively (Zopfi et al., 2008). Samples for H_2_S (monobromobimane) as well as filtered samples for NO_3_^-^ concentration and nitrate isotope analyses were stored frozen. Samples for NO_2_^-^ and NH_4_^+^ concentration analysis were stored at 4 °C. If samples contained significant amounts of nitrite, aliquots were treated with sulfamic acid to remove nitrite. For incubation experiments with natural microbial consortia, we filled unfiltered lake water from selected water depths into 1-L sterile borosilicate glass bottles. The bottles were stored at 4 °C before adding substrates and starting the incubations within 5 days. DNA for phylogenetic analysis (from same depth as water for incubations) was extracted by filling sterile 1-L borosilicate bottles with lake water. Within ∼12 h of sampling, we filtered ∼1.2 L through 0.2 µm cyclopore track edged membrane filters (Whatman). Filters were frozen immediately and stored at -70 °C.

### Lake La Cruz

Lake La Cruz is a small karstic lake (∼132 m diameter) in Central-Eastern Spain, close to the city of Cuenca, at an altitude of ∼1000 m above sea level. It has a depth of ∼20 m (Camacho et al., 2017), and has been meromictic for more than 300 years, with a chemocline ∼5 m above the lake bottom and hypolimnetic waters rich in dissolved ions such as ferrous iron, bicarbonate, and NH_4_^+^. We collected water samples during sampling campaigns in March 2015 and March 2017 at the deepest point of the lake (39°59’20’’N, 01°52’25’’W) at the RTZ. We used an in situ peristaltic pump with gas-tight tubing to fill sterile 0.5 or 2-L borosilicate bottles with water from 14.5 m (2015 and 2017) and 17 m (2017) for incubations. In addition, we extracted DNA from water samples collected at 14.5 m in 2017, as described for Lake Lugano.

### 2.3. Lake water incubation experiments

We performed incubation experiments with Lake Lugano water samples (“Exp_LL”) from November 2016, February 2017, October 2017, and April 2018. The procedure and concentration data are presented in Tischer et al. (2025). Briefly, we applied an N_2_ headspace to samples in 1-L borosilicate bottles and purged for 30 minutes with N_2_ before adding substrates (final concentration ∼25 µM for NO_3_^-^ and acetate and ∼25 to 100 µM for H_2_S) from sterile and anoxic stock solutions (5 - 10 mM KNO_3_, 5 - 10 mM Na_2_S·9H_2_O, 5 - 10 mM C_2_H_3_NaO_2_) for three main treatments (in duplicates): NO_3_^-^, NO_3_^-^ + acetate, and NO_3_^-^ + H_2_S. The bottles were incubated in the dark at ∼8 °C. At each of the 3 to 16 time points, we collected ∼20 ml of water through the septum using a syringe and a sterile canula for NO_3_^-^ and NO_2_^-^ concentration and NO_3_^-^ isotope analysis, as described in section 2.1. When NO_3_^-^ and NO_2_^-^ were completely consumed (between 3 and 25 days), or the NO_3_^-^ concentrations did not significantly change, the experiments were terminated, and the remaining water was filtered for DNA analysis. Lake La Cruz incubation experiments (“Exp_LC”) were executed following principally the same procedure, using 0.5-L borosilicate bottles with water from 14.5 m depth (2015 and 2017), and 250 ml serum vials, closed with butyl rubber stoppers and aluminum seals, filled with water from 17 m depth (2017). Lake La Cruz incubations were performed at room temperature in the dark.

### 2.4. Concentration measurements and isotope analysis

#### Concentrations of N and S compounds

We determined concentrations of NO_x_ by reducing it to NO in acidic vanadium (V^3+^), followed by chemiluminescent detection of NO (Antek Model 745; Braman and Hendrix 1989). NO_2_^-^ concentrations were measured photometrically (Hansen and Koroleff, 1999), and NO_3_^-^ concentrations were then calculated as the difference of NO_x_ and NO_2_^-^ concentrations. In addition, we used nitrate concentration data of the entire water column of Lake Lugano obtained by a monitoring program coordinated by the University of Applied Sciences and Arts of Southern Switzerland (www.cipais.org). Concentrations of NH_4_^+^ and H_2_S were determined using common photometric methods (Cline, 1969; Krom, 1980). Samples for H_2_S (bimane method) were analyzed using high-performance liquid chromatography (HPLC, Dionex), followed by fluorescence detection (Zopfi et al., 2008). The concentration data from Lake Lugano, except for February, August, and December 2015, were previously presented (Tischer et al., 2025).

### Nitrate isotope ratios

We analyzed NO_3_^-^ isotope ratios using the denitrifier method (Sigman et al., 2001). Briefly, 20 nmoles of sample NO_3_^-^ were converted to N_2_O by cultured denitrifying bacteria (*Pseudomonas aureofaciens*). Subsequently, the N_2_O was automatically extracted, purified, and analyzed using a customized purge-and-trap system coupled to a continuous-flow GC-IRMS (Thermo Scientific DELTA V Plus). The NO_3_^-^ isotope measurements were calibrated against international NO_3_^-^ standards IAEA-N3 (δ^15^N = 4.7‰, δ^18^O = 25.6) and USGS34 (δ^15^N = 1.8‰, δ^18^O = 27.9‰), and an internal standard (UBS-1; δ^15^N = 14.15‰, δ^18^O = 25.7‰). A selection of samples was measured in duplicate, and the reproducibility for δ^15^N was <0.3‰ and for δ^18^O <0.4‰.

### 2.5. Turnover rate and N and O isotope effect calculation

The NO_3_^-^ reduction rates in incubations were calculated based on the decrease in NO_3_^-^ concentration over time, including samples showing changes in concentration with available isotope measurements. NO_2_^-^ production rates were calculated similarly for pure culture experiments. Both NO_3_^-^ reduction and NO_2_^-^ production rates are reported with positive values. We estimated the isotope effects of N (^15^ε) and O (^15^ε) imparted on nitrate during dissimilative nitrate reduction in incubation experiments and in situ by fitting δ^15^N and δ^18^O values of NO_3_^-^ to the linear equations of Mariotti et al. (1981):

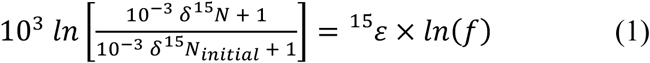

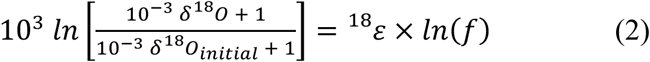

where *f* represents the remaining fraction of the substrate NO_3_^-^, i.e. [NO_3_^-^]/[NO_3_^-^]_initial_. For the incubation experiments, [NO_3_^-^]_initial_ and δ^15^N_initial_ (δ^18^O_initial_) represent the initial nitrate pool at t = 0, whereas for the application of the closed system "Rayleigh model" to nitrate data in the Lake Lugano water column, [NO_3_^-^]_initial_ and δ^15^N_initial_ (δ^18^O_initial_) were taken as the nitrate concentration and isotopic composition of the subsurface hypolimnion above the RTZ. Isotope values at low [NO_3_^-^] that strongly deviated from a linear relationship in a Rayleigh plot were removed when calculating the isotope effects. Graphical presentations of the nitrate concentration versus isotope ratio relationship were made by plotting δ^15^N or δ^18^O against the natural logarithm of *f*, where the negative slope of the linear regression is a good approximation of ^15^ε and ^18^ε, respectively (Mariotti et al., 1981). Here, we report normal kinetic isotope effects, which reflect higher reaction rates for the lighter isotopologues, as positive values. The nitrate Δδ^18^O:Δδ^15^N ratio was calculated by performing a linear regression analysis between δ^18^O and δ^15^N. The regression analysis generally resulted in R^2^ values of >0.90. For Inc_LL and Inc_LC, we calculated isotope effects considering both duplicates, yet we excluded experiments, where less than 40% of the nitrate was consumed.

### 2.6. Statistical analysis

We performed two-tailed *t*-tests for all isotope values and rates to test whether they were significantly different. Statistical analyses of Δδ^18^O:Δδ^15^N, ^15^ε, and ^18^ε determined through (1) culture and (2) lake water incubations were conducted in R (Version 1.4.1103). For each isotope effect, an initial analysis of covariance (ANCOVA) with categorical and continuous variables was performed using the function aov(). For the pure culture incubations, we used ’nitrate reductase’, ‘carbon source’, ’initial NO_3_^-^ concentration’, ’NO_3_^-^ reduction rate’, and ’NO_2_^-^ production rate’ as explaining factors and variables, respectively. For the lake water incubations, we analyzed ’treatments’ (i.e., added substrates), ’lakes’ (Lake Lugano versus Lake La Cruz), the interaction between ’treatments’ and ’lakes’, and ’NO_3_^-^ reduction rate’ by using values of each replicate where >40% NO_3_^-^ was consumed. Subsequently, we applied a stepwise backward regression using the Bayesian Information Criterion (BIC) (Schwarz, 1978; Burnham and Anderson, 2002) using the step() function in R. The BIC is defined as:

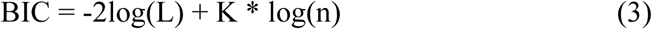

where L is the maximized value of the empirical likelihood function, K is the number of estimable parameters, and n is the sample size. The procedure removes response variables and factors until the most likely model with the lowest BIC remains. The resulting factors and/or variables were used in final ANOVA (analysis of variance) or ANCOVA models (function aov()). In addition, we performed post-hoc tests using Tukey’s honest significance test with the TukeyHSD() function (Haynes, 2013) to test for significance between groups or using regressions with the dependent variable and the covariate. We used the same approach to verify the potential influence of the environmental variables ’NO_3_^-^ flux’, ’NH_4_^+^ flux’, ’H_2_S flux’ (data from Tischer et al., 2025), ’primary productivity’ (data from www.cipais.org), ’depth of O_2_ depletion’, and/or ’season’ on the in situ nitrate N and O isotope effects in the Lake Lugano water column. We used the R package ggplot2 (Wickham, 2016) for graphical representations.

### 2.7. Phylogenetic analysis

We extracted DNA from in situ and incubation samples using the FastDNA Spin Kit for Soil (MP Biomedicals). Amplicons for Illumina sequencing were generated by PCR using the universal primers 515F-Y and 926R, targeting the V4 and V5 regions of the 16S rRNA gene (Parada et al., 2016). The procedure of library preparation, sequencing, and data processing, including the generation of amplicon sequence variants (ASVs) by denoising amplicons to zero-radius operational taxonomic units, has been described in detail (Su et al., 2023). The taxonomic assignment of ASVs was done using the SILVA reference database v138. Further analyses were done using R Version 4.1.1 including the phyloseq package version 1.36.0 (McMurdie and Holmes, 2013). Data were transformed into relative abundances (RA) of total counts by dividing the read counts of a specific ASV by the total reads per sample. Based on the change in the relative abundance (ΔRA), we identified the most responsive taxa in one duplicate incubation in some of the lake water incubation experiments (“Exp_LL” and “Exp LC”), and we determined whether they harbor *narG* and/or *napA*, based on their genome or genomic information of close relatives, using the Kyoto Encyclopedia of Genes and Genomes (www.kegg.jp).

## 3. Results

### 3.1. Nitrate N and O isotope effects associated with denitrification in pure cultures

In the incubation experiments with pure cultures of *Pseudomonas* sp. and *E. coli*, NO_3_^-^ was completely consumed within 5 to 100 h, accompanied by the transient accumulation of NO_2_^-^ (Fig. S1). Nitrate reduction by *Pseudomonas* sp. strains (Exp_culture_P1 and Exp_culture_P2) yielded Rayleigh isotope effects (Table 1, Fig. S2) in a wide range between 7.9‰ and 30.2‰ for ^15^ε, and between 8.5 and 28.2 for ^18^ε. While the ^15^ε and ^18^ε differed between experiments and strains, the corresponding Δδ^18^O:Δδ^15^N ratios were robust with values close to 1 (between 0.92 and 0.99). The *E. coli* strains (Exp_culture_Ecoli) reduced NO_3_^-^ with N and O isotope effects in a much a narrower and lower range between 8.0 to 16.7‰ for ^15^ε and 5.9 to 16.4‰ for ^18^ε. Nitrate reduction by strain JCB4011 (*narG*) produced an average Δδ^18^O:Δδ^15^N of 0.98, while strain JCB4031 (*napA*) yielded significantly lower Δδ^18^O:Δδ^15^N trajectories of 0.65 to 0.74.

The statistical analysis (Table 2, Fig. S3) showed much unexplained variance (42.5%) of the observed ^15^ε values. Across the different pure-culture incubations, only the carbon source could explain some of the variance yet without statistical significance (ANOVA; F = 3.8, R^2^ = 0.58, *p* = 0.05). Similarly, ^15^ε did not differ systematically among strains with a different type of reductase. In contrast, for ^18^ε, 55.9% of the variance of could be explained by the type of nitrate reductase, with significant differences between the *narG* + *napA* strains (20.6‰ ± 1.8‰ standard error, SE) and the *napA* strain (6.2‰ ± 0.3‰ SE) (ANOVA; F = 7.6, R^2^ = 0.56, *p* = 0.007). Observed differences in Δδ^18^O:Δδ^15^N ratios were almost exclusively explained by the type of nitrate reductase (ANOVA; F = 70.0, R^2^ = 0.90, *p* < 0.001). That is, the average Δδ^18^O:Δδ^15^N for nitrate reduction by cultures with the *napA* strain (0.70 ± 0.05 SE) was clearly distinct from that for cultures possessing the *narG* strain (0.97 ± 0.01 SE) or the *narG*+*napA* strains (0.95 ± 0.01 SE) (Tukey post-hoc test; *p* < 0.001).

**Table 2.**
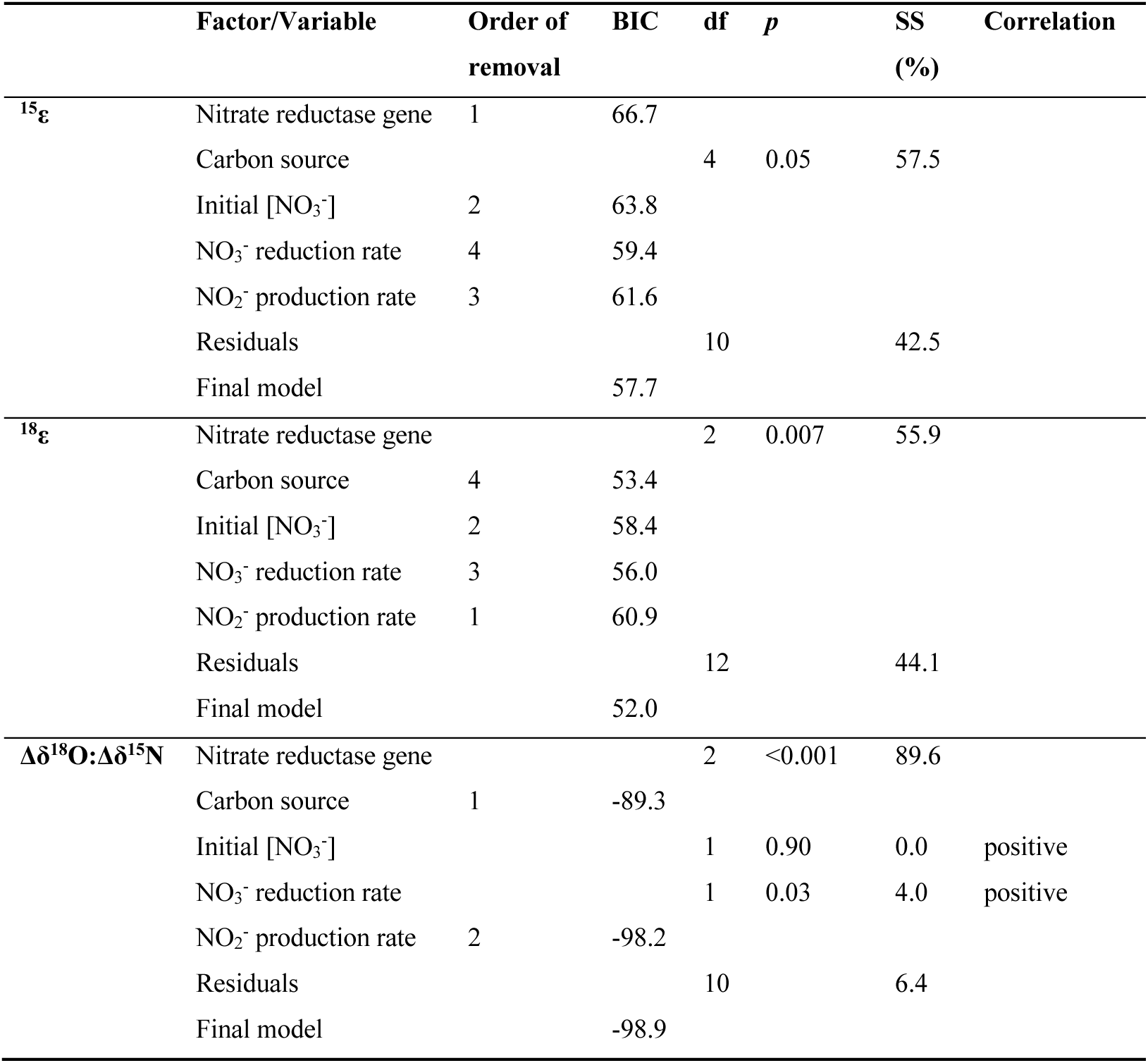
Factors and variables explaining ^15^ε, ^18^ε, and Δδ^18^O:Δδ^15^N in the pure culture incubation experiments with strains of *Pseudomonas* sp. and *Escherichia coli*. The initial model considers all potentially relevant factors and variables. The final model determined via the Bayesian Information Criterion (BIC), selects for the most important parameters. The *p* values (statistical significance given for *p* < 0.05), degrees of freedom (df), sum of squares (SS), and type of correlation between the dependent variable and the covariate (positive/negative, for continuous variables) are provided for the remaining traits in the final model. See section 2.6. for details.

| | Factor/Variable | Order of removal | BIC | df | $p$ | SS (%) | Correlation |
| --- | --- | --- | --- | --- | --- | --- | --- |
| $^{15}\epsilon$ | Nitrate reductase gene | 1 | 66.7 | | | | |
|  | Carbon source |  |  | 4 | 0.05 | 57.5 |  |
| | Initial $[\text{NO}_3^-]$ | 2 | 63.8 | | | | |
| | $\text{NO}_3^-$ reduction rate | 4 | 59.4 | | | | |
| | $\text{NO}_2^-$ production rate | 3 | 61.6 | | | | |
|  | Residuals |  |  | 10 |  | 42.5 |  |
|  | Final model |  | 57.7 |  |  |  |  |
| $^{18}\epsilon$ | Nitrate reductase gene | | | 2 | 0.007 | 55.9 | |
|  | Carbon source | 4 | 53.4 |  |  |  |  |
| | Initial $[\text{NO}_3^-]$ | 2 | 58.4 | | | | |
| | $\text{NO}_3^-$ reduction rate | 3 | 56.0 | | | | |
| | $\text{NO}_2^-$ production rate | 1 | 60.9 | | | | |
|  | Residuals |  |  | 12 |  | 44.1 |  |
|  | Final model |  | 52.0 |  |  |  |  |
| $\Delta\delta^{18}\text{O}:\Delta\delta^{15}\text{N}$ | Nitrate reductase gene | | | 2 | <0.001 | 89.6 | |
|  | Carbon source | 1 | -89.3 |  |  |  |  |
| | Initial $[\text{NO}_3^-]$ | | | 1 | 0.90 | 0.0 | positive |
| | $\text{NO}_3^-$ reduction rate | | | 1 | 0.03 | 4.0 | positive |
| | $\text{NO}_2^-$ production rate | 2 | -98.2 | | | | |
|  | Residuals |  |  | 10 |  | 6.4 |  |
|  | Final model |  | -98.9 |  |  |  |  |

### 3.2. Incubation experiments with natural consortia

In most Lake Lugano incubations (Exp_LL) with only NO_3_^-^ added, nitrate was not fully consumed, levelling off at about 5-30% of the initial amount. In contrast, in all incubations where acetate or H_2_S was added in addition to NO_3_^-^ (Fig. S4a-c; data adopted from Tischer et al., 2025), NO_3_^-^ was completely consumed. In the Lake La Cruz incubations, both in the NO_3_^-^ -only and the NO_3_^-^ + H_2_S treatment, all NO_3_^-^ was reduced (Fig. S5a-b). In all incubations, NO_2_^-^ was produced and accumulated permanently or transiently (Fig. S4d-f, Fig. S5c-d). δ^15^N and δ^18^O of NO_3_^-^ systematically increased with decreasing substrate concentrations in all incubations. Considering only those incubations exhibiting fractional NO_3_^-^ consumption of >40%, ^15^ε values ranged between 13 and 30‰, and ^18^ε values between 9 and 26‰, giving rise to Δδ^18^O:Δδ^15^N ratios ranging between 0.58 and 1.04 (Table 3, Fig. 1, Fig. S6).

**Figure 1.**
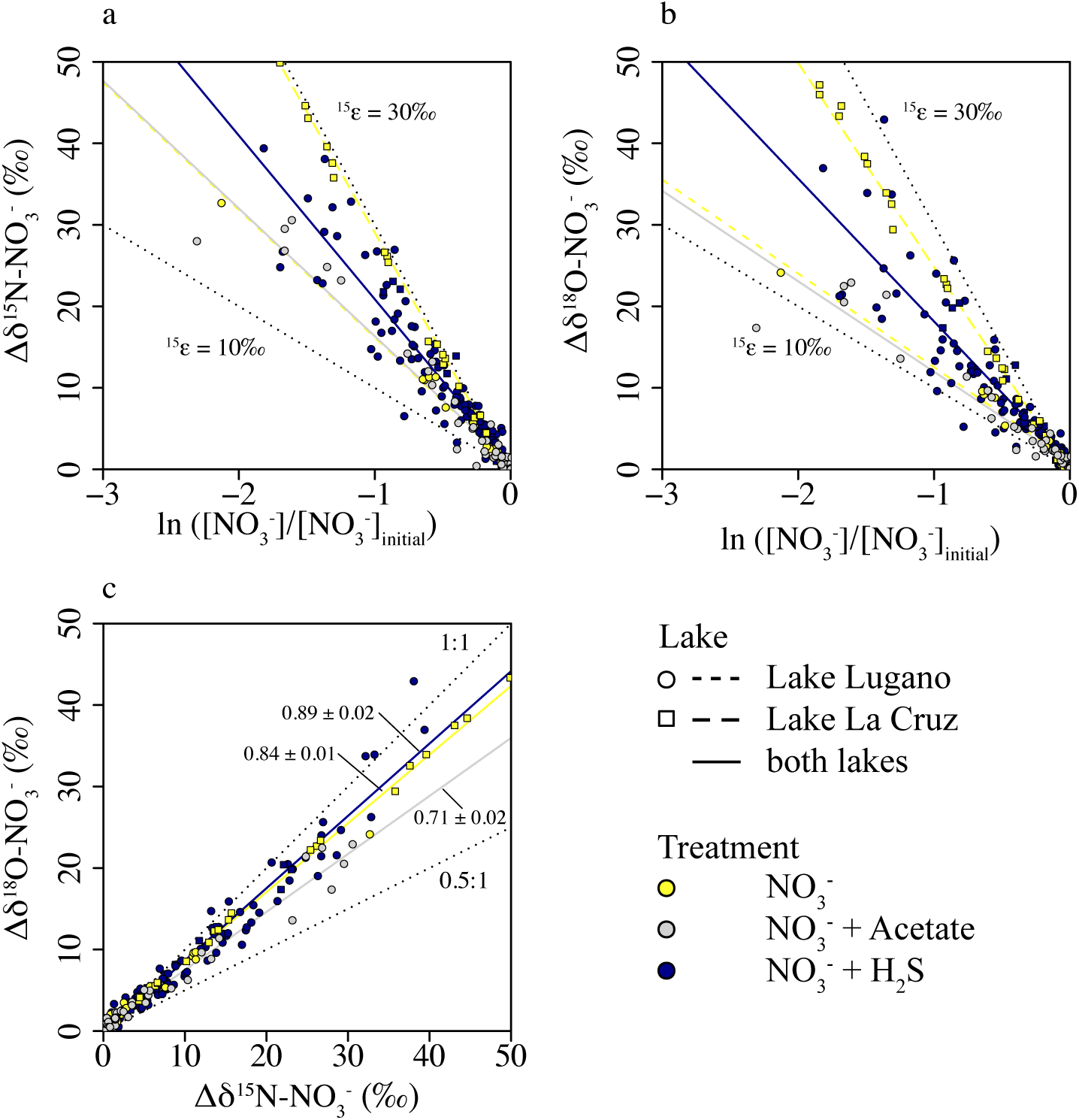
**a)** δ^15^N-Rayleigh-plot, **b)** δ^18^O-Rayleigh plot, and **c)** Δδ^18^O-vs-Δδ^15^N plot for nitrate isotope data for incubation experiments with natural microbial consortia from the Lake Lugano and the Lake La Cruz water column, amended with NO_3_^-^, NO_3_^-^ + acetate (Lake Lugano only), or NO_3_^-^ + H_2_S.

**Table 3.** Nitrate isotope fractionation for nitrogen (^15^ε) and oxygen (^18^ε), and Δδ^18^O:Δδ^15^N ratios determined in incubation experiments using water from Lake Lugano (Exp_LL) or Lake La Cruz (Exp_LC). Data are presented with standard error of means. Non-significant results are indicated by “ns”.

| Lake | Time of Sampling | Depth | Initial [NO<sub>3</sub><sup>-</sup>] (μM) | Additional substrate | $^{15}\epsilon$ (‰) | $^{18}\epsilon$ (‰) | $\Delta\delta^{18}\text{O}:\Delta\delta^{15}\text{N}$ | NO<sub>3</sub><sup>-</sup> reduction (μmol L<sup>-1</sup> h<sup>-1</sup>) | |
| --- | --- | --- | --- | --- | --- | --- | --- | --- | --- |
|  |  |  |  |  |  |  |  | Duplicate 1 | Duplicate 2 |
| Lugano | November 2016 | 95 m | 25 |  | 17.7 ± 1.1 | 13.8 ± 1.6 | 0.77 ± 0.08 | 2.6 ± 0.5 <sup>a</sup> |  |
|  |  |  | 25 | 25 μM acetate | 18.4 ± 0.8 | 14.4 ± 0.7 | 0.78 ± 0.03 | 5.6 ± 2.4 (ns) | 4.6 ± 1.1 |
|  |  |  | 25 | 25 μM H<sub>2</sub>S | 23.4 ± 1.4 | 23.6 ± 2.1 | 1.04 ± 0.04 | 1.2 ± 0.2 | 0.8 ± 0.1 |
| Lugano | November 2016 | 105 m | 25 | 25 μM H<sub>2</sub>S | 24.2 ± 1.4 | 18.8 ± 1.2 | 0.58 ± 0.02 | 0.9 ± 0.0 | 0.9 ± 0.1 |
| Lugano | November 2016 | 155 m | 25 | 25 μM acetate | 16.9 ± 1.0 | 10.4 ± 0.4 | 0.61 ± 0.01 | 1.9 ± 0.7 | 1.6 ± 0.5 |
|  |  |  | 25 | 25 μM H<sub>2</sub>S | 20.6 ± 1.3 | 19.1 ± 1.4 | 0.93 ± 0.03 | 0.9 ± 0.1 | 0.8 ± 0.1 |
| Lugano | February 2017 | 95 m | 25 | 25 μM acetate | 19.5 ± 1.5 | 14.8 ± 1.0 | 0.75 ± 0.03 | 1.6 ± 0.5 | 1.1 ± 0.5 (ns) |
|  |  |  | 25 | 70 μM H<sub>2</sub>S | 25.5 ± 1.1 | 23.4 ± 0.9 | 0.90 ± 0.04 | 0.5 ± 0.1 | 0.8 ± 0.1 |
| Lugano | October 2017 | 105 m | 25 | 25 μM acetate | 17.4 ± 1.0 | 11.8 ± 0.6 | 0.68 ± 0.01 | 2.2 ± 0.6 | 1.5 ± 0.7 (ns) |
|  |  |  | 25 | 30 μM H<sub>2</sub>S | 19.9 ± 1.7 | 16.9 ± 1.6 | 0.86 ± 0.03 | 2.4 ± 0.5 | 1.9 ± 0.1 |
| Lugano | April 2018 | 105 m | 25 |  | 15.1 ± 0.3 | 11.1 ± 0.3 | 0.74 ± 0.01 | 5.1 ± 2.3 (ns) <sup>a</sup> |  |
|  |  |  | 25 | 25 μM acetate | 13.4 ± 1.0 | 9.1 ± 1.1 | 0.69 ± 0.05 | 4.7 ± 1.5 | 4.3 ± 2.8 (ns) |
|  |  |  | 25 | 100 μM H<sub>2</sub>S | 14.6 ± 0.8 | 12.1 ± 0.7 | 0.82 ± 0.02 | 2.5 ± 0.3 | 2.4 ± 0.2 |
| La Cruz | March 2015 | 14.5 m | 100 |  | 27.1 ± 0.2 | 22.1 ± 0.2 | 0.82 ± 0.00 | 1.0 ± 0.1 | 0.8 ± 0.1 |
| La Cruz | March 2017 | 14.5 m | 100 |  | 30.4 ± 0.8 | 25.9 ± 0.6 | 0.85 ± 0.01 | 1.2 ± 0.1 | 1.2 ± 0.1 |
|  |  |  | 100 | 250 μM H<sub>2</sub>S | 25.4 ± 1.7 | 21.1 ± 1.9 | 0.84 ± 0.03 | 4.6 ± 0.6 | 4.5 ± 0.4 |
| La Cruz | March 2017 | 17 m | 25 |  | 29.0 ± 0.7 | 25.4 ± 0.4 | 0.87 ± 0.01 | 2.1 ± 0.1 <sup>a</sup> |  |
|  |  |  | 25 | 250 μM H<sub>2</sub>S | 27.0 ± 0.2 | 25.1 ± 0.3 | 0.93 ± 0.01 | 2.0 ± 0.7 (ns) <sup>a</sup> |  |

<sup>a</sup> In only one of the duplicate experiments >40% of the initial NO<sub>3</sub><sup>-</sup> was consumed.

The final ANCOVA models (Table 4) for ^15^ε and ^18^ε values showed that the values depended significantly on the lake (^15^ε: F = 48.8, R^2^ = 0.52, *p* < 0.001; ^18^ε: F = 18.9, R^2^ = 0.32, *p* < 0.001) but also on the NO_3_^-^ reduction rate (^15^ε: F = 17.7, R^2^ = 0.21, *p* < 0.001; ^18^ε: F = 11.5, R^2^ = 0.22, *p* < 0.001) (Fig. S7a-d). In particular, observed ^15^ε and ^18^ε values were higher in experiments with natural microbial consortia from Lake La Cruz (Exp_LC; ^15^ε = 27.9‰ ± 0.8‰ SE, ^18^ε = 23.7‰ ± 0.9‰ SE) than in experiments with consortia from Lake Lugano (Exp_LL; ^15^ε = 19.5‰ ± 0.8‰ SE, ^18^ε = 16.2‰ ± 1.1‰ SE). As for the latter, ^15^ε and ^18^ε decreased significantly with increasing nitrate reduction rate (linear regression; ^15^ε: -1.7 ± 0.4‰ per 1 µmol NO_3_^-^ L^-1^ d^-1^, R^2^ = 0.46, *p* < 0.001; ^18^ε: -2.1 ± 0.6‰ per 1 µmol NO_3_^-^ L^-1^ d^-1^, R^2^ = 0.37, *p* = 0.002). Δδ^18^O:Δδ^15^N values changed with treatment (ANOVA; F = 9.5, R^2^ = 0.41, *p* < 0.001; Table 4), with significant differences between the NO_3_^-^ + acetate (Δδ^18^O:Δδ^15^N = 0.72 ± 0.03 SE) and the NO_3_^-^ + H_2_S treatment (Δδ^18^O:Δδ^15^N = 0.89 ± 0.03 SE) (Tukey post-hoc test; *p* < 0.001), whereas the NO_3_^-^ treatment (Δδ^18^O:Δδ^15^N = 0.82 ± 0.02 SE) was not significantly different from the other treatments (Table 5, Fig. S7e).

**Table 4.**
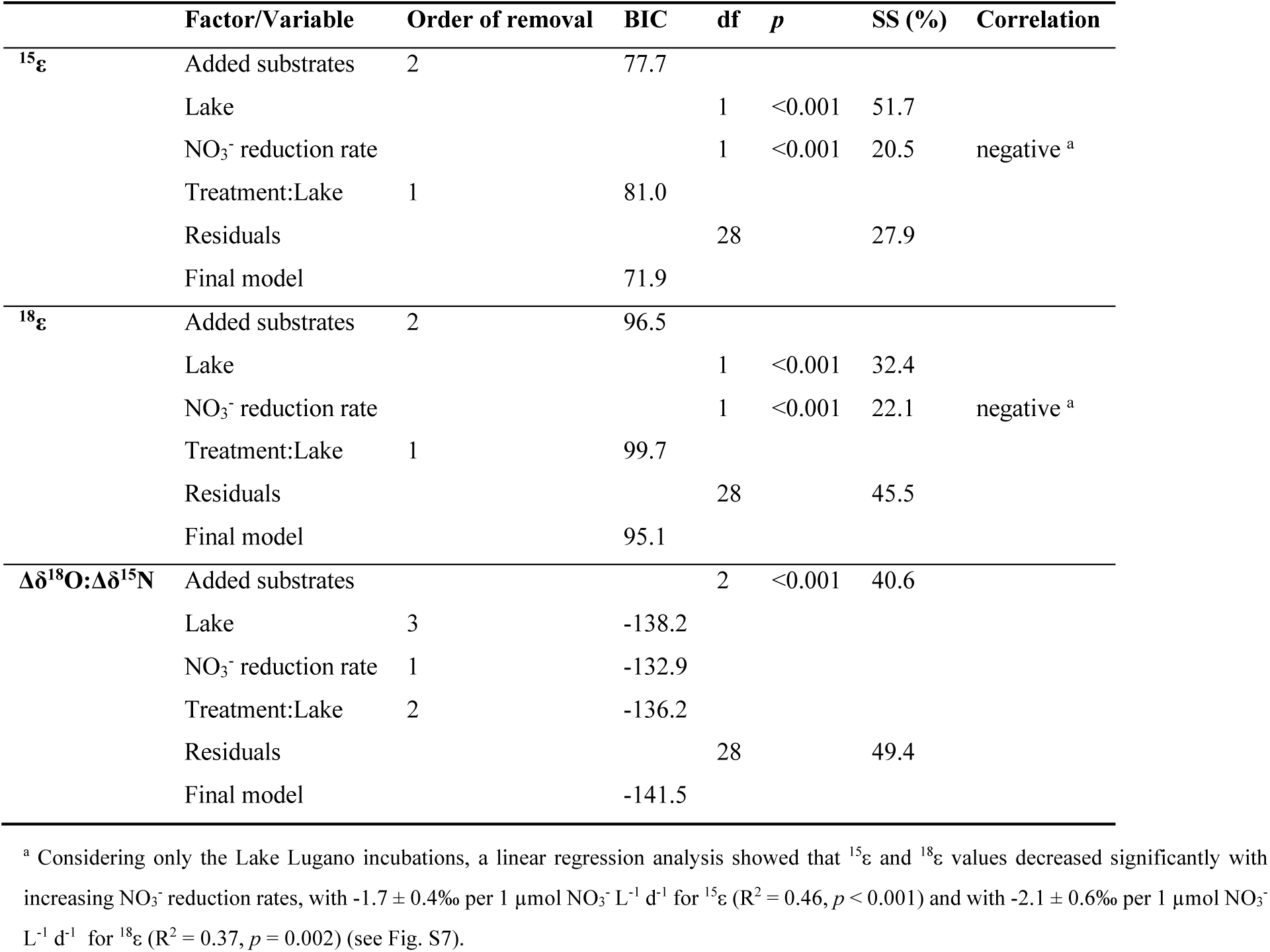
Factors and variables explaining ^15^ε, ^18^ε, and Δδ^18^O:Δδ^15^N in the incubation experiments with lake water from Lake Lugano and Lake La Cruz. The initial model considers all potentially relevant factors and variables. The final model, determined via the Bayesian Information Criterion (BIC), selects for the most important parameters. The *p* values (statistical significance given for *p* < 0.05), degrees of freedom (df), sum of squares (SS), and type of correlation between the dependent variable and the covariate (positive/negative, for continuous variables) are provided for the remaining traits in the final model. See section 2.6 for details.

**Table 5.**
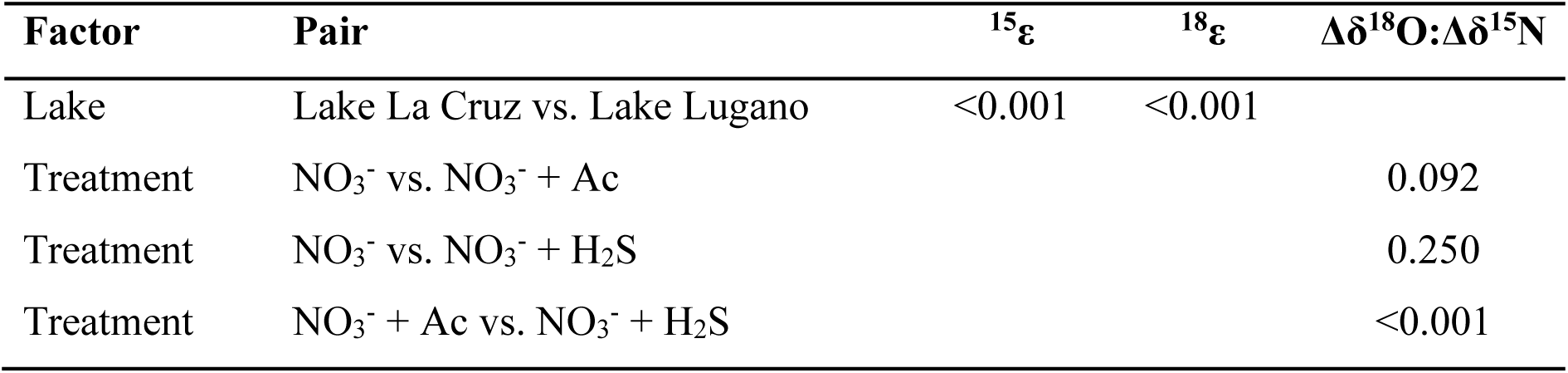
Tukey post hoc tests *p* values for significant differences within groups retained in the final statistical models explaining ^15^ε, ^18^ε and Δδ^18^O:Δδ^15^N selected by Bayesian Information Criterion (Table 4).

The analysis of 16S rRNA genes (Fig. 2) revealed that in most EXP_LL incubations, *Dechloromonas* sp. (sequenced representatives harboring *napA*) was the most competitive taxon, with large increases in relative abundance (ΔRA = 9 - 75%) over the course of the incubations. Enrichment of this taxon was highest in incubations with NO_3_^-^ + acetate, and other N-reducing taxa were not enriched at all during the incubation. In the NO_3_^-^ + H_2_S amended treatments, different taxa got enriched, including predominantly sulfur-dependent nitrate-reducing bacteria such as *Sulfuricurvum* sp. (*napA*), *Desulfomonile* sp. (*narG*+*napA*), *Hydrogenophaga* sp. (*narG*), and *Thiobacillus* sp. (*narG*). During Exp_LC incubations, the potentially N-reducing organisms *Hydrogenophaga* sp., *Nitrospira* sp. (*narG*+*napA*), and *Rhodoferax* sp. (*narG*) were enriched, with ΔRA between 2 and 11%.

**Figure 2.**
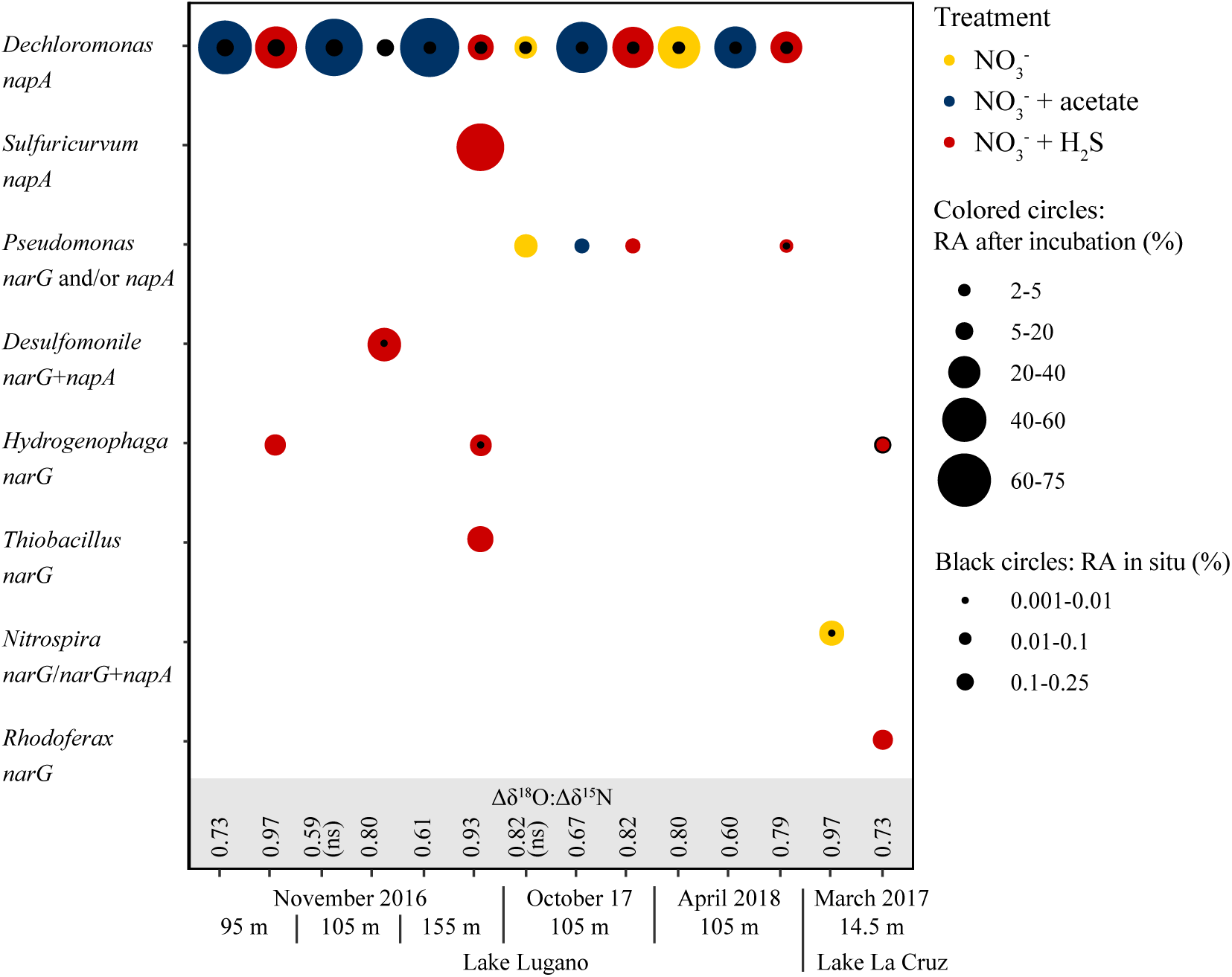
Relative abundance (RA) of bacterial genera harboring nitrate reductase genes (*narG* and/or *napA*) that enriched in incubation experiments with natural microbial consortia from Lake Lugano or Lake La Cruz, amended with NO_3_^-^, NO_3_^-^ + acetate (Lake Lugano only), or NO_3_^-^ + H_2_S. The Δδ^18^O:Δδ^15^N ratios determined in the same incubation experiment are shown. Non-significant values are indicated by “ns”. For details see section 2.7.

### 3.3. Observations in the water column of Lake Lugano

At the sampling timepoints, the RTZ in the Lake Lugano North Basin was mostly located between ∼85 and 125 m (Fig. S8). Nitrate reached concentrations up to 35 µM in the upper oxic water column and was completely consumed within the RTZ (Fig. 3a, Fig. S8). NO_2_^-^ concentrations in the RTZ were mostly below 0.05 µM, with few exceptions. For example, in April 2015 and April 2018, maximum NO_2_^-^ concentrations within the RTZ reached 0.5 µM and 0.16 µM, respectively. NH_4_^+^ began to accumulate within the RTZ, whereas H_2_S concentrations started to increase at the bottom of the RTZ (Fig. S8).

**Figure 3.**
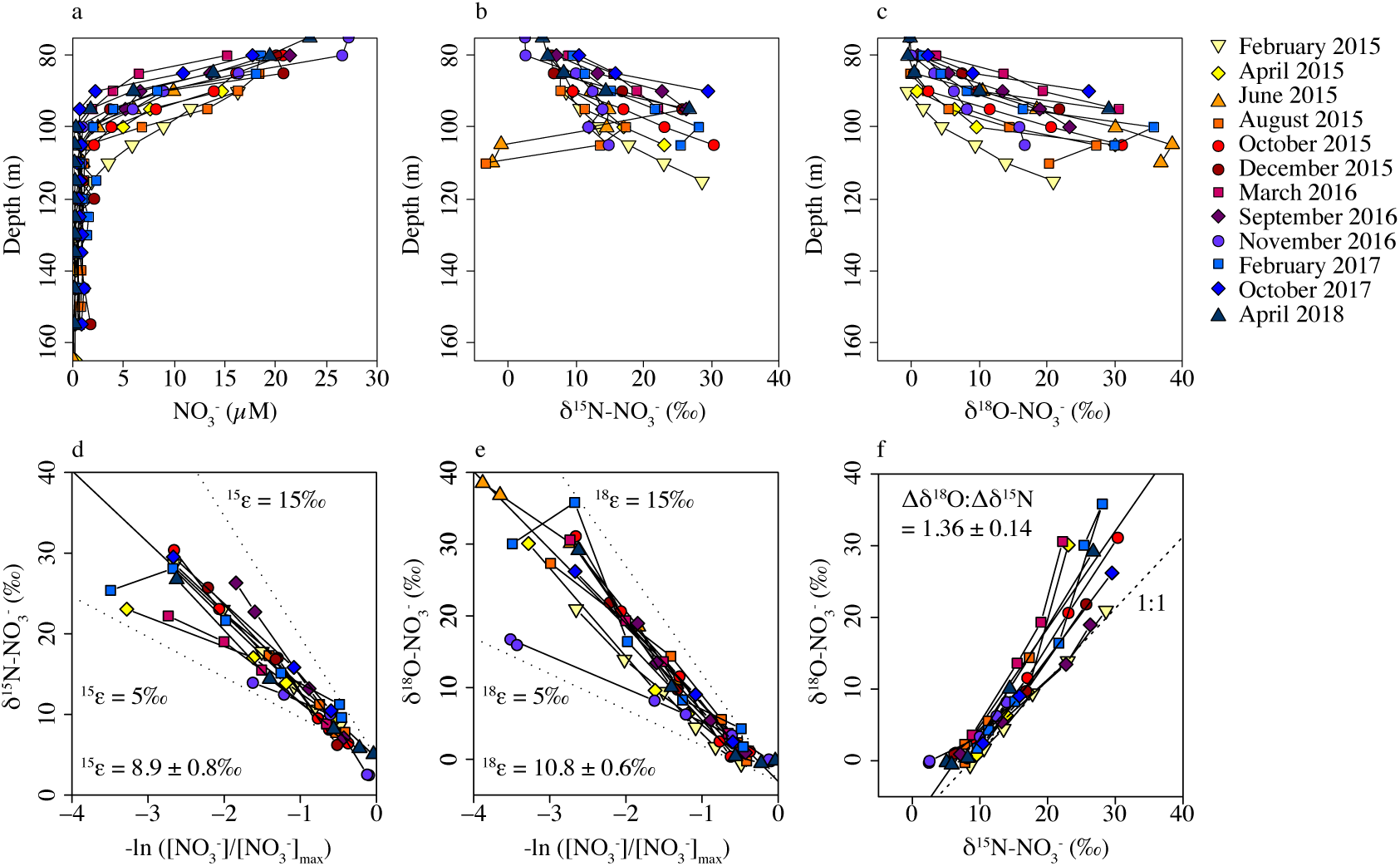
Nitrate concentrations and isotope ratios between 75 m and 165 m in the water column of the Lake Lugano North Basin. **a)** NO_3_^-^ concentrations, **b)** δ^15^N-NO_3_^-^ values, **c)** δ^18^O-NO_3_^-^ values, **d)** δ^15^N-Rayleigh-plot, **e)** δ^18^O-Rayleigh plot, and **f)** δ^18^O-vs-δ^15^N plot. For panels d - f, values strongly deviating from a 1:1 relationship were excluded. The slope of the regression line, representing the average of the sampling campaigns, is given with the standard error of means.

The δ^15^N-NO_3_^-^ values above and within the RTZ increased with decreasing nitrate concentrations from ∼5 - 10‰ around 80 m to values up to 30‰ near the depth of complete NO_3_^-^ depletion (Fig. 3b), indicating N-isotope fractionation with progressing nitrate consumption. Corresponding to the δ^15^N-NO_3_^-^ trend (in the same part of the water column, for the different samplings, respectively), δ^18^O-NO_3_^-^ increased from ∼0‰ to up to 40‰ (Fig. 3c). In some of the nitrate isotope profiles, δ^15^N and δ^18^O values decreased with the samples of lowest NO_3_^-^ concentration, as observed in June and August 2015, where δ^15^N values dropped to <0‰, and the nitrate δ^18^O decreased by 2 - 8‰ (Fig. S8). Rayleigh model-based ^15^ε values ranged between 4.8‰ and 13.5‰ (average 8.9 ± 0.8‰ SE), very similar to the ^18^ε value range between 4.8 and 13.8‰ (average 10.8 ± 0.6‰ SE) (Fig. 3d-e). Δδ^18^O:Δδ^15^N ratios for the different water column nitrate isotope profiles ranged between 0.67 and 2.15, with an average of 1.36 ± 0.14 SE (Table S2, Fig. 3f). We did not observe any significant effects of seasonally variable hydrochemical parameters on ^15^ε, ^18^ε, or Δδ^18^O:Δδ^15^N values (data not shown).

### 4. Discussion

In the following sections, we discuss the controls on the Rayleigh N and O nitrate isotope effects during nitrate reduction in incubation experiments, and the implications for the generation of nitrate dual isotope signatures in lakes, including enzymatic effects and the influence of reaction rates and substrate limitation. Since the observed isotope effects exhibited a large unexplained variability, we will, in the second part of the discussion, focus on the much more robust dual O versus N isotope enrichment ratio. Analyzing the results of the incubation experiments with pure cultures and natural microbial consortia from the two different lakes, we discuss the factors potentially affecting Δδ^18^O:Δδ^15^N ratios, such as the type of nitrate-reducing organism/metabolism involved (i.e., organotrophic versus lithotrophic nitrate reduction), as well as the type of enzyme that catalyzes nitrate reduction. In addition, turning to the water column nitrate isotope data from Lake Lugano, we address how isotopic overprinting by simultaneous nitrate production and a high N turnover in a net-denitrifying zone may shape the coupled nitrate O:N isotope signature in the natural environment. Finally, we critically evaluate the potential of nitrate isotopes in general to identify N-transformation processes in aquatic environments.

### 4.1. Controls on Rayleigh nitrate isotope fractionation effects in incubation experiments and in situ

#### 4.1.1. High unexplained variation in Rayleigh isotope effects

The wide range of nitrate N and O isotope effects observed both in the incubation experiments with pure cultures and with natural microbial consortia (^15^ε = 12 - 32 ‰, ^18^ε = 8 - 31 ‰), as well as in situ (^15^ε = 5 - 14 ‰, ^18^ε = 5 - 14 ‰), remains largely unexplained, making Rayleigh modeled isotope effects a difficult tool for tracing nitrate reduction mechanisms and their controls. In this study, we found that substrate and diffusion limitation (see section 4.1.2.), as well as nitrate reduction rates (see section 4.1.3.) impacted Rayleigh isotope effects, but not the carbon source or nitrite production rate. We did not test for other potential influencing factors, though, such as pH and cell-specific nitrate reduction rates. In general, the Rayleigh N and O isotope effects associated with nitrate reduction can vary significantly, depending on various cellular and/or enzymatic mechanisms (Granger et al., 2008). For example, Nar is located at the inner cell membrane, with the active site facing the cytoplasm, requiring active NO_3_^-^ transport across the inner membrane. Nap, on the other hand, is located in the periplasm, and relies on passive diffusion of NO_3_^-^ through porins in the outer cell membrane. Diffusive NO_3_^-^ transport is generally regarded as reversible and thought to impart only negligible isotope fractionation. The first irreversible step of NO_3_^-^ reduction is the enzymatic cleavage of the N-O bond, resulting in the enrichment of the residual nitrate in the heavy isotopes ^15^N and ^18^O (Granger et al., 2008; Treibergs and Granger, 2017). However, whether the enzymatic N and O isotope effects are propagated to the external nitrate pool (i.e., observable either in the lake water column or in the medium of an incubation experiment), depends on the extent to which intracellular ^15^N- and ^18^O-enriched nitrate is transported out of the bacterial cell (Nar) and diffuses through the outer membrane (Nar and Nap). For Nar, the ratio of active nitrate uptake into the cell versus nitrate efflux has been suggested to be the driving variable altering the intrinsic (enzymatic) isotope effects expressed in the NO_3_^-^ pool outside the cell (Shearer et al., 1991; Granger et al., 2008; Kritee et al., 2012). Variability in ^15^ε and ^18^ε for Nap likely depends on limitation of nitrate supply, which in turn depends on Michaelis-Menten substrate kinetics of Nap activity, cell-specific nitrate reduction rates, and/or the supply rates of nitrate (Granger et al., 2008; Frey et al., 2014b).

The high variability in N and O isotope effects reported here is in line with observations made in multiple other culture and field studies (^15^ε = 5 - 33‰, ^18^ε = 4 - 33‰, Table S1), where incubation conditions could at least partly explain variations in isotope effects (Delwiche and Steyn, 1970; Mariotti et al., 1981; Granger et al., 2008; Frey et al., 2014b). In comparison, pure-enzyme assays of nitrate reductases exhibit much less variable N and O isotope effects (Karsh et al., 2012; Treibergs and Granger, 2017). Some of the isotope effect variability observed in our study could be explained by variation in the cell-specific nitrate reduction rates (i.e., nitrate consumption rates divided by cell numbers), which have been shown to correlate positively with ^15^ε (Kritee et al., 2012; Frey et al., 2014b).

#### 4.1.2. Influence of substrate and diffusion limitation

Naturally expressed isotope effects in the Lake Lugano water column, both in 2009-2010 (Wenk et al., 2014) and in 2015-2018 (this study), are much lower compared to in vitro experiments with natural consortia (Exp_LL; Fig. 4a). In contrast to denitrification in the batch experiments, where closed-system Rayleigh dynamics apply, denitrification in the natural environment often exhibits reduced isotope fractionation. This reduction in the apparent isotope effect may be partly due to the fact that strict “Rayleigh model rules” are to some extent violated when modeling nitrate isotope signatures in the natural spatial heterogeneity of a water column. That is, nitrate is not necessarily drawn from a homogenous pool, but denitrification occurs primarily where the nitrate pool has already been pre-enriched in ^15^N by partial consumption (Deutsch et al., 2004; Altabet, 2007; Wenk et al., 2014).

**Figure 4.**
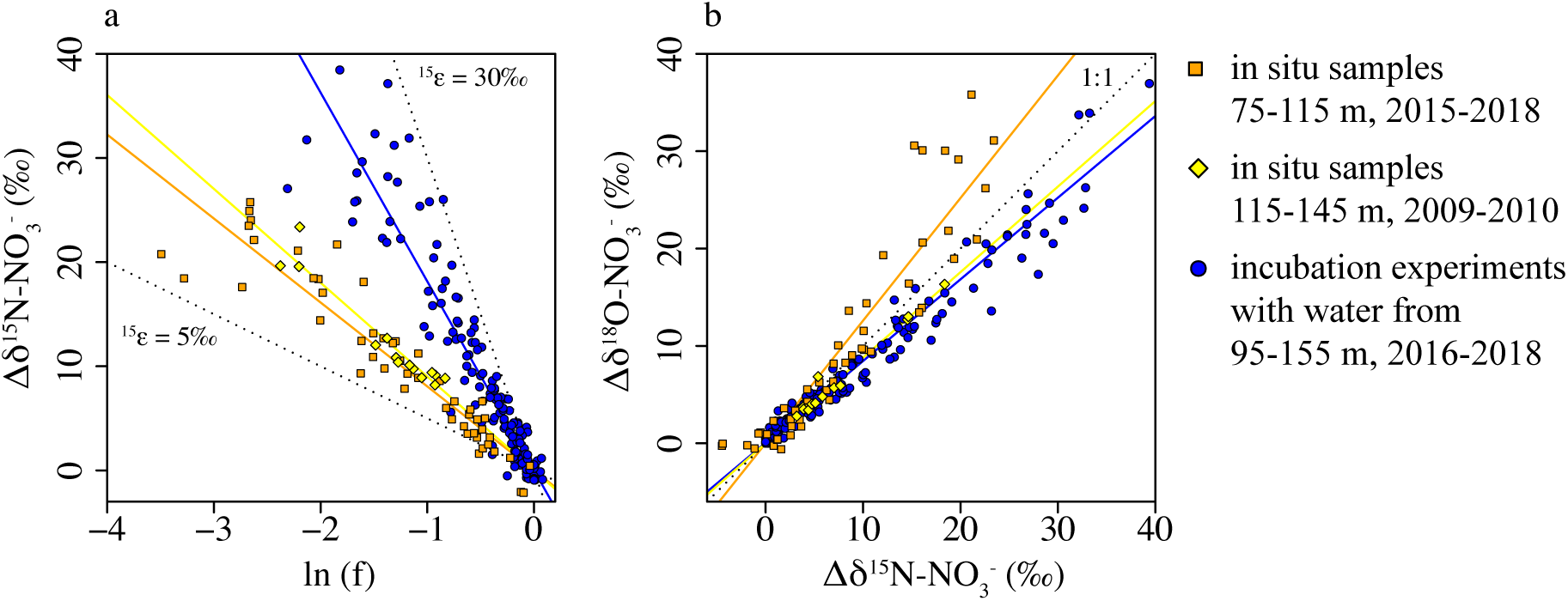
**a)** δ^15^N-Rayleigh-plot and **b)** Δδ^18^O:Δδ^15^N plot for nitrate in situ samples from the Lake Lugano North Basin and from incubation experiments with natural microbial consortia from Lake Lugano.

Moreover, open system aspects (i.e., diffusive resupply and/or nitrate production; see section 4.1.3.) may apply, lowering the net ecosystem N isotope effect for water column denitrification. In agreement with previous work (Wenk et al., 2014), we argue however that, beyond the potential bias when applying the Rayleigh approach to the water column data, the ecosystem nitrate isotope effect observed here is indeed relatively low. A likely explanation for the decrease in ^15^ε and ^18^ε values is the restricted substrate availability when utilization is strong, and supply is limited by diffusion (at the organism level). Such limitation can decrease the commitment to the enzymatic reaction, reducing the isotope fractionation (Kritee et al., 2012; Treibergs and Granger, 2017). Indeed, restricted availability of the electron acceptor NO_3_^-^ and/or the most likely electron donors (H_2_S or organic compounds) is evident from the observation that the active denitrifying zone in the North Basin is essentially devoid of H_2_S and low in NO_3_^-^ (<5 µM) (Tischer et al., 2025). The in situ NO_3_^-^ concentrations are almost certainly much lower than the Michaelis-Menten constant (K_m_) values for nitrate uptake and for the enzymatic activity of the dissimilatory nitrate reductases (3 µM to the mM range; Sparacino-Watkins et al., 2014), which likely explains the small ecosystem-scale Rayleigh isotope effects compared to those observed in the incubation experiments, where both NO_3_^-^ and electron donors were available at much higher concentrations.

#### 4.1.3. Influence of nitrate reduction rates

Even when substrate limitation is not evident, high nitrate reduction rates may lead to diffusion limitation at the organism/enzyme level and may thus lead to reduced ^15^ε and ^18^ε values. Indeed, low ^15^ε and ^18^ε appeared to correspond to increased NO_3_^-^ reduction rates, as shown in Exp_LL (Fig. S7a+c). Similar observations have been made in soil incubation experiments (Mariotti et al., 1982) and culture studies (Bryan et al., 1983) testing the isotope effects imparted by denitrification. In contrast, in Exp_LC and in other culture and field studies (Granger et al., 2008; Snider et al., 2009), no systematic correlations between denitrification rates and isotope effects are observed. We speculate that the differing observations between Lake La Cruz and Lake Lugano are due to microbial community differences and varying nitrate affinities. Lake La Cruz supports much higher in situ denitrification rates (up to 987 nmol N-N_2_ L^-1^ d^-1^; Tischer et al., 2022) than Lake Lugano (up to 113 nmol N-N_2_ L^-1^ d^-1^; Tischer et al., 2025), suggesting higher organic carbon availability. Despite this difference, we observed similar nitrate reduction rates in incubations (Table 3), showing that the stimulation of denitrification was more pronounced in Lake Lugano. Furthermore, Exp_LL was dominated by a single organism (*Dechloromonas* sp.), while in Exp_LC we could not observe the enrichment of any dominant species, indicating that, here, active resident denitrifiers were already capable of reducing the added nitrate in Lake La Cruz water. As a consequence, there was a lower cell density of denitrifiers at the starting point of Exp_LL compared to Exp_LC, ultimately leading to diffusion limitation in Exp_LL but not in Exp_LC. This highlights that a negative correlation between NO_3_^-^ reduction rates and ^15^ε values cannot always be expected, and suggests that certain conditions may be required. For instance, in an environment with limited availability of NO_3_^-^ (and electron donors), the sudden stimulation of nitrate reduction by availability of NO_3_^-^ (e.g., >10 µM) (and an electron donor such as organic carbon or H_2_S) can result in the rapid growth and activity of a denitrifying community.

In the water column of Lake Lugano, we do not necessarily expect any significant lowering of ^15^ε values due to high NO_3_^-^ reduction rates, since NO_3_^-^ concentrations were low in the active denitrifying layer. In addition, observed ^15^ε values in the water column (8.9 ± 0.8‰ SE) were generally much lower compared to the incubations of Exp_LL (21.2‰ ± 1.2‰ SE) and hence, the effect of substrate limitation probably dominated over any possible modulating effects by variable nitrate reduction rates.

### 4.2. Role of Nar and Nap activity on the dual nitrate O vs N isotope signature in incubation experiments

#### 4.2.1. Dual nitrate isotope effects in pure culture experiments

A mechanistic difference between the two nitrate reductases with regards to coupled nitrate N and O isotope fractionation is clearly indicated by the different Δδ^18^O:Δδ^15^N ratios for Nar (∼0.8 - 1.0) compared to Nap (∼0.5 - 0.6) observed in the pure culture experiments. The significantly lower Δδ^18^O:Δδ^15^N ratios in association with Nap activity, for example by *E. coli* strain JCB4031, can be primarily attributed to the comparatively low ^18^ε. Also in previous culture studies, observed ^18^ε values did not exceed 18‰ for nitrate-reducing organisms with *napA* only, while denitrifiers possessing *narG* or both *narG* and *napA* expressed ^18^ε values up to 33‰ (Table S1). The exact biochemical mechanism explaining the lower ^18^ε for Nap than for Nar remains uncertain, but as discussed in Frey et al. (2014b) and Treibergs and Granger (2017), it might be related to different substrate binding to the molybdenum (Mo) center of the active site of the different enzymes, respectively. The Mo center is an integral part of both Nar and Nap, but despite the fact that the two molybdoenzymes ultimately catalyze the same biochemical reaction, they may bind and cleave nitrate molecules differently, possibly leading to a less pronounced enrichment of ^18^O_NO3_ (i.e. lower ^18^ε) in the remaining nitrate pool in the Nap-mediated reduction, as compared to an equivalently high ^18^O and ^15^N enrichment in the Nar-mediated reduction. Independent of the underlying catalytic mechanisms of the two nitrate reductases, and despite a certain (unexplained) variability even for pure culture studies (with Δδ^18^O:Δδ^15^N values of 0.80 - 1.04 for *narG* and 0.50 - 0.68 for *napA*; Table S1), the Δδ^18^O:Δδ^15^N signature provides a more robust signal than the corresponding individual N and O isotope effects, and may serve as an excellent diagnostic tool to detect activities of Nar versus Nap.

#### 4.2.2. Impact of microbial players and metabolisms in natural consortia

Based on our findings from incubations involving natural microbial consortia from Lake Lugano and Lake La Cruz, we provide putative evidence that varying substrate conditions affect enzymatic activity and the Δδ^18^O:Δδ^15^N ratios by selectively stimulating different microbial players/communities. We did not quantify *narG* and *napA* genes or transcripts in the incubation samples. Hence, we speculate that the observed experimental Δδ^18^O:Δδ^15^N ratio range between 0.58 and 1.04 largely reflects differences in the relative activity of Nar versus Nap, and not, for example, changes in the relative contribution of nitrate reduction versus production (in contrast to the water column; see section 4.3.). More specifically, in these incubations, given the anoxic conditions, we can exclude the overprinting effect of nitrification. Moreover, based on most recent results demonstrating negligible anammox rates in both the Lake Lugano and Lake La Cruz water columns (Tischer et al., 2022, 2025), we assume that the impact of nitrate production by anammox is small.

In the NO_3_^-^-amended experiments without additional electron donor, we observed intermediate Δδ^18^O:Δδ^15^N slopes (0.81 ± 0.02 SE), which we assume to represent a realistic isotopic baseline of nitrate reduction in situ. The values lie between the average endmember values for Nar and Nap (0.9 - 1.0 and 0.5 - 0.7, respectively), reflecting the involvement of a natural community of microbes that harbors both *narG* and *napA*. For Lake Lugano, we previously showed that a suite of different N-reducing bacteria co-exists in the water column, including *Halomonas* sp. (*narG or napA*), *Candidatus* Accumulibacter sp. (*napA*), and *Denitratisoma* sp. (*narG* and *napA*) (Tischer et al., 2025; www.kegg.jp). Compared to the NO_3_^-^-only control treatment, the addition of acetate significantly reduced the Δδ^18^O:Δδ^15^N ratio. This is most likely due to the activation of organotrophic NO_3_^-^ reduction, particularly via Nap, as indicated by the dominant proliferation of *napA-*harboring *Dechloromonas* sp. (Salinero et al., 2009), previously identified as an important denitrifier in a nitrate-polluted aquifer (Duffner et al., 2021). In contrast, the addition of H_2_S did not cause a significant shift in the Δδ^18^O:Δδ^15^N ratio relative to the nitrate-only control. This is surprising, as S-dependent denitrification has previously been associated with Nap (Frey et al., 2014b). However, the S-oxidizers identified in Lake Lugano are known to possess *narG* and/or *napA*, including *Thiobacillus* sp. *(narG*), *Sulfuricurcum* sp. (*napA*), *Sulfurimonas* sp. (*napA*), or *Sulfuritalea* sp. (*narG* and *napA*) (Tischer et al., 2025). Indeed, NO_3_^-^ + H_2_S additions in both Exp_LL and Exp_LC led most often to a seemingly balanced enrichment of *narG* and/or *napA*-harboring taxa, including *Hydrogenophaga* sp. *(narG*), *Sulfuricurcum* sp. (*napA*), or *Desulfomonile* sp. (*narG* and *napA*), such that in the end the observed Δδ^18^O:Δδ^15^N ratios (0.90 ± 0.03 SE) were similar to the Δδ^18^O:Δδ^15^N in the nitrate-only control/baseline experiment. Still, single experiments with added H_2_S resulted in ratios up to 0.97, even when *Dechloromonas* sp. was enriched (Fig. 2). Hence, not all of the variability in Δδ^18^O:Δδ^15^N in the incubations with natural denitrifying consortia can be explained by the relative partitioning of the active microorganisms reducing nitrate with either Nar or Nap. Nevertheless, the natural-consortia incubations with Lake Lugano and Lake La Cruz samples, and the corresponding isotope and phylogenetic analyses, clearly show that Nar and Nap enzyme activity can be equally important in freshwater environments, likely explaining the Δδ^18^O:Δδ^15^N ratios <1 in the studied lakes. What exactly controls the relative importance of Nar versus Nap, and how reliably Δδ^18^O:Δδ^15^N ratios can inform about the nitrate reductase partitioning and the underlying catalytic mechanisms should be a focus of future research.

#### 4.2.3. Impact of C/N ratios on the relative activity of Nar versus Nap

In organisms possessing both *narG* and *napA*, the ratio of electron donor (organic C) to electron accepter (e.g., nitrate), and corresponding C and N substrate limitations, respectively, seem to control the activity of Nar and Nap, which is indicated in our incubation experiments with pure cultures and natural consortia. Under conditions with high [NO_3_^-^], catalysis by Nar seems to be preferred over Nap, as seen, for example, in the incubations with *Pseudomonas* sp. strains ELC2NRS8 and ELC1RS10 (media with >200 µM NO_3_^-^), where the Δδ^18^O:Δδ^15^N signal was constantly ≥ 0.90 (Table 1). This observation aligns with previous research indicating the preeminence of Nar over Nap under elevated NO_3_^-^ concentrations (Potter et al., 1999; Asamoto et al., 2021). Moreover, the *E. coli* strain employing Nar demonstrated markedly faster nitrate reduction rates compared to the strain using Nap under equivalent high [NO_3_^-^] conditions, suggesting that Nar is advantageous for energy conservation. On the other hand, higher electron donor/electron acceptor ratios confer a selective advantage to Nap over Nar, as recently documented in studies on nitrate reduction by *P. aeruginosa* and by *Pseudogulbenkiania* sp., leading to lower Δδ^18^O:Δδ^15^N values of 0.50 to 0.73 (Chen et al., 2020; Asamoto et al., 2021).

### 4.3. Dual nitrate O vs N isotope effects in the water column of the Lake Lugano North Basin

#### 4.3.1. Isotopic overprinting by nitrification

Co-occurring contribution of newly nitrified nitrate to the nitrate pool near the RTZ of the Lake Lugano North Basin is evidenced by significant positive deviations of δ^18^O:δ^15^N-NO_3_^-^ slopes from a 1:1 ratio (1.36 ± 0.14 SE), and inflections at low [NO_3_^-^]. The high Δδ^18^O:Δδ^15^N ratios are in accordance with modelled δ^18^O-to-δ^15^N trajectories that Granger and Wankel (2016) predicted for conditions where denitrification and nitrification occur simultaneously (at a relatively high nitrite oxidoreductase (Nxr)/Nar ratio), yet, under conditions with barely any O-atom exchange between nitrite and water. Indeed, in the Lake Lugano RTZ, we rarely observed the accumulation of NO_2_^-^. In addition, we previously provided evidence that the RTZ is an active cryptic N-cycling zone including denitrification and nitrification (Tischer et al., 2025). The greater enrichment in ^18^O-NO_3_^-^ compared to ^15^N-NO_3_^-^ can be attributed to the branching isotope effect during the reduction of NO_3_^-^ to NO_2_^-^ (and subsequently to N_2_O), which adds to, and to some extent counteracts, the normal O isotope effects imparted by Nar/Nap (and Nir); this isotope branching involves the preferential abstraction of ^16^O from the N-O bond, producing NO_2_^-^ with elevated δ^18^O relative to the nitrate from which it formed (Casciotti et al., 2002, 2007). Upon (re-)oxidation of the nitrite, this high-δ^18^O signal is then reflected in the δ^18^O of the NO_3_^-^ pool, which is concurrently reduced and produced (yet the latter with higher δ^18^O values relative to the δ^15^N values; Granger and Wankel 2016). In contrast, O-atom exchange between NO_2_^-^ and H_2_O (with a δ^18^O_H2O_ of ∼-7‰ for the Lake Lugano water column; Lehmann et al., 2003), would act to lower the δ^18^O of NO_3_^-^ relative to the δ^15^N of NO_3_^-^, leading to Δδ^18^O:Δδ^15^N values <1 (Granger and Wankel, 2016). However, such O-atom exchange is only expected, if nitrite accumulates and remains in the ambient water for several days (Casciotti et al., 2007; Buchwald and Casciotti, 2013), which is generally not the case in Lake Lugano. In this regard, the Δδ^18^O:Δδ^15^N values >1 appear to rule out such scenario, and rather indicate the superimposed effect of NO_3_^-^ production via nitrite oxidation.

Apart from Δδ^18^O:Δδ^15^N ratios above unity, we argue that also the positive inflections of the Δδ^18^O:Δδ^15^N trajectory in the low [NO_3_^-^] range (Fig. 3f) can be seen as indicator of active nitrification in the water column. Our observed nitrate isotope trends differ from the model results of Granger and Wankel (2016). In their idealized model simulations, in which, for a given model scenario, a constant Nxr/Nar ratio was assumed, the Δδ^18^O:Δδ^15^N trajectory flattened at higher isotope values because the δ^18^O of newly nitrified NO_3_^-^ increased less relative to the δ^18^O of the decreasing residual NO_3_^-^ pool. In Lake Lugano, the relative rates of nitrification and denitrification likely change over depth, giving rise to shifts in the apparent δ^18^O vs δ^15^N trends that differ from those predicted by Granger and Wankel (2016). That is, nitrification rates are likely highest around the RTZ, where O_2_ and NH_4_^+^ concentration profiles overlap. At shallower depths above the RTZ, NO_3_^-^ concentrations are higher and the isotopic signal is more influenced by diffusion and mixing with nitrate from the overlying oxic water (Wenk et al., 2014). In contrast, at low [NO_3_^-^], δ^15^N-NO_3_^-^ values are particularly prone to the impact of oxidation of ^15^N-depleted NO_2_^-^ produced from NH_4_^+^. With δ^15^N-NH_4_^+^ values below the RTZ of 8 - 10‰ (Wenk et al., 2014) and assuming an average isotope effect of Δδ^15^N_NH4_^+^◊_NO2_^-^ ≈ 30‰ (Denk et al., 2017), the δ^15^N of steady-state NO_2_^-^ production from NH_4_^+^ oxidation should fall around -20‰ (Fig. 5a). On the other hand, the δ^15^N of NO_2_^-^ produced from NO_3_^-^ is with ∼ 16‰ much higher, assuming δ^15^N-NO_3_^-^ values in the RTZ of ∼30‰ (Fig. 3b) and an average Nar isotope effect of Δδ^15^N_NO3_^-^◊_NO2_^-^ ≈ 14‰ (Denk et al., 2017) (Fig. 5b).

**Figure 5.**
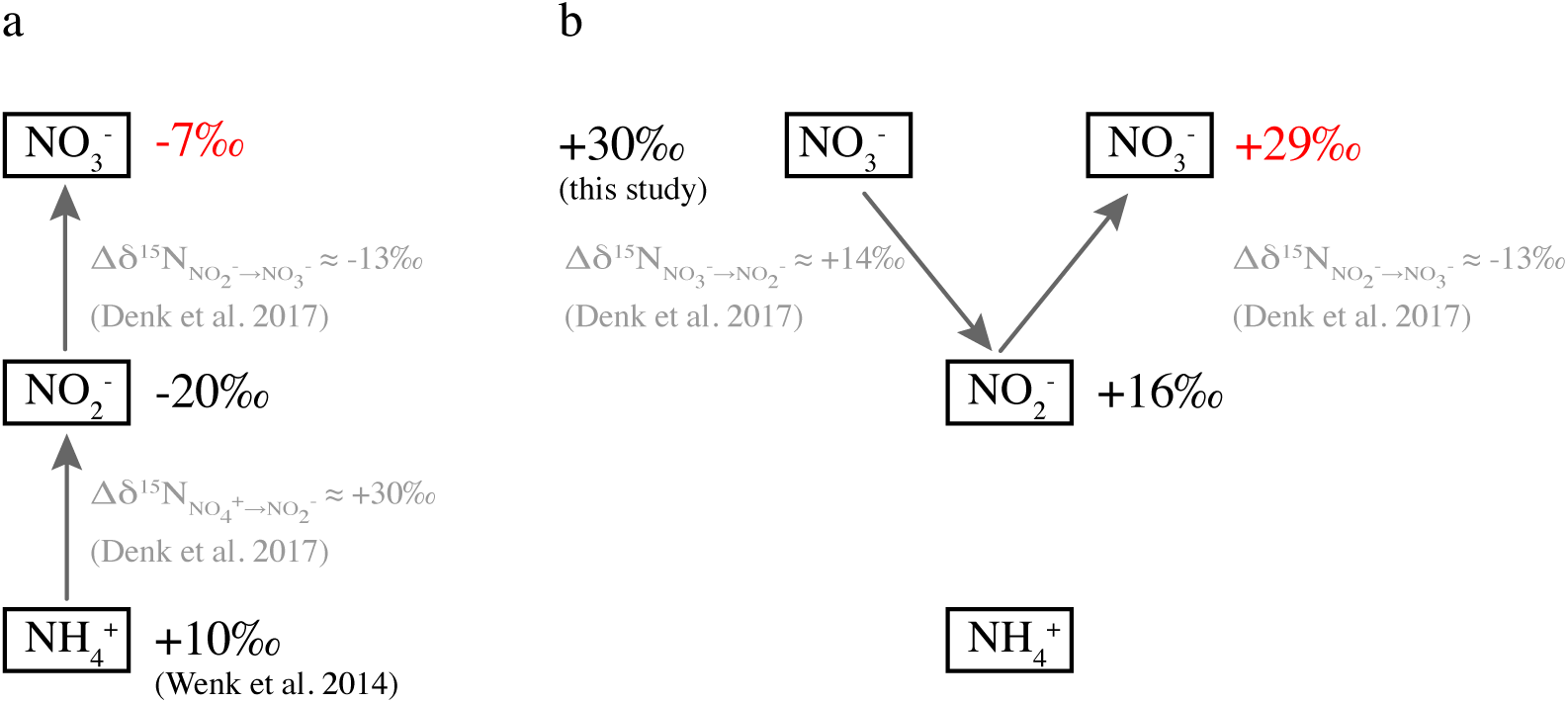
Isotopic composition of nitrate produced during nitrification in the redox transition zone of the Lake Lugasyno North Basin depending on the origin of the precursor nitrite. **a)** Scenario, where NO_3_^-^ is produced via ammonia oxidation and nitrite oxidation. **b)** Scenario, where dissimilatory NO_3_^-^ reduction is coupled with NO_2_^-^ re-oxidation. Isotope effect associated to ammonia oxidation, nitrite oxidation, and nitrate reduction are average values from Denk et al. (2017).

Therefore, the inverse isotope effect of NO_2_^-^ re-oxidation to NO_3_^-^ of Δδ^15^N_NO2_^-^◊_NO3_^-^ = -13‰ (Casciotti, 2009) will result in low δ^15^N-NO_3_^-^ of ∼-7‰ for nitrified NO_3_^-^ while coupled NO_3_^-^reduction and production produces ^15^N signatures close to the initial δ^15^N-NO_3_^-^ with ∼29‰. Indeed, we found δ^15^N-NO_3_^-^ values as low as -2 to -3‰ in June and August 2015, suggesting almost 100% of nitrate directly originating from NH_4_^+^ oxidation.

The positive curvature (i.e., upward bending) of the Δδ^18^O:Δδ^15^N slopes corresponding to the low-nitrate concentration region of the water column may in parts also be attributed to increasing δ^18^O-NO_3_^-^ (at NO_3_^-^ concentrations ∼3 - 10 µM). Fractional O_2_ consumption enriches the residual O_2_ pool in ^18^O, so that particularly at low O_2_ concentrations, high δ^18^O_2_ (>30 ‰) can be expected (Mader et al., 2017). As a consequence, within the RTZ (i.e., in the low-nitrate portion of the water column), NO_2_^-^ produced from the oxidation of NH_4_^+^, where one O is incorporated from O_2_ (Casciotti et al., 2010), is reflected by elevated δ^18^O-NO_3_^-^ values of the nitrified NO_3_^-^ (e.g., in February 2017 or April 2018), increasing the nitrate O-vs-N isotope decoupling towards higher Δδ^18^O:Δδ^15^N ratios at low-nitrate concentrations in most of the nitrate isotope profiles. While we cannot fully comprehend, why, in the same context, the δ^18^O-NO_3_^-^ values sporadically drop at NO_3_^-^ <3 µM paralleling δ^15^N-NO_3_^-^ values (Fig. S8), we nevertheless argue that the generally positive inflections of the δ^18^O:δ^15^N slopes are a good indicator for an increasing influence of nitrite oxidation activity.

The relatively high in situ Δδ^18^O:Δδ^15^N ratios with up to 2.15 ± 0.45 SE, and the inflection of the Δδ^18^O:Δδ^15^N slopes stand in contrast to the isotopic Δδ^18^O:Δδ^15^N-NO_3_^-^ baseline of 0.81 ± 0.02 SE determined in anoxic experiments with natural microbial consortia from the same lake basin (Exp_LL). The high Δδ^18^O:Δδ^15^N for the in situ measurements seem exceptional for a freshwater ecosystem (see Table S1), and it remains uncertain as to what exactly causes these high values. However, the discrepancy to the observations in the natural-consortia experiments is explained by the overprinting effect of nitrate production in the natural water column, which does not occur under the anoxic conditions in the incubation experiments. Hence, the results presented here underscore that, in particular under low [NO_3_^-^] conditions in freshwater systems, dual nitrate isotope signatures can provide an excellent means to qualitatively diagnose and pinpoint high nitrification rates and N recycling in natural waters.

#### 4.3.2. Temporal variation of nitrate dual isotope effects

Based on the discussion in section 4.3.1., the relatively large range in Δδ^18^O:Δδ^15^N values between 0.67 and 2.15 observed for different time points in the Lake Lugano water column suggests temporal changes in the ratio of nitrate reduction and production and/or in the relative importance of Nar and Nap. A variable importance of individual N-transforming processes, yet without a detected seasonal dependence, is supported by observed changes in measured N-transformation rates (e.g., denitrification rates ranging between 29 to 113 nmol N-N_2_ L^-1^ d^-1^) and by variable abundances of N-cycling genera (Tischer et al., 2025). For example, 16S rRNA data show maximum relative abundances in the RTZ of the Lake Lugano North Basin from 2015 to 2018 that vary between 0.9 to 3.0% for the dominant nitrite oxidizer *Nitrospira* sp. and between 0.1 and 1.8% for one of the main identified nitrate reducers *Dechloromonas* sp. Given the meromictic character of the Lake Lugano North Basin, with its stable chemocline at relatively great depth (Tischer et al., 2025), the cause for the observed temporal changes remains uncertain. We suggest that periods of elevated O_2_ concentrations and occasional O_2_ intrusions into anoxic waters, respectively, stimulate the activity of aerobic microorganisms as ammonia and nitrite oxidizers and inhibit anaerobic nitrate reduction. The biweekly monitoring of O_2_ concentrations in the water column confirms that the penetration depth of O_2_ into the hypolimnion can change within a relatively short period of time (Tischer et al., 2025; www.cipais.org), hence modulating the conditions that regulate redox processes.

Interestingly, Wenk et al. (2014) reported a relatively constant ratio of Δδ^18^O:Δδ^15^N ratio of 0.89 ± 0.05 around the RTZ in the same lake basin in 2009-2010 (Fig. 4b), clearly lower than the observed values of 1.36 ± 0.14 in this study. On the one hand, their data indicate that over the study period of 6 months, the conditions were more or less stable. In comparison, we detected differences in Δδ^18^O:Δδ^15^N ratio of up to 1.1 within only a few months. On the other hand, the discrepancy between the two datasets suggests that in the O_2_-deficient water layer of the Lake Lugano North Basin, the influence of nitrification on the nitrate budget has become much more pronounced relative to nitrate reduction. The mechanisms behind the potential rise in the significance of nitrate regeneration by nitrification are not fully understood. However, it could be linked to the overall transition of the water column following the last mixing events in 2005/2006. As suggested by Su et al. (2023), during this period slow-growing nitrifying taxa may have required time to establish a stable population.

#### 4.3.3. Implications for marine environments

Our results clearly indicate that both Nap activity and isotopic overprinting can cause Δδ^18^O:Δδ^15^N trajectories of NO_3_^-^ deviating from a 1:1 ratio, contrary to previous assumptions. In marine denitrifying waters, where Nap has traditionally been considered subordinate or negligible, Δδ^18^O:Δδ^15^N ratios greater than 1:1 have been attributed to isotopic overprinting by nitrification. However, our results suggest that some of these systems may have been characterized by a nitrate isotopic baseline of <1 due to the contribution from Nap activity, but with isotopic overprinting by NO_3_^-^ production, pushing the Δδ^18^O:Δδ^15^N ratios >1. This may be particularly true for the Baltic Sea, where a δ^18^O:δ^15^N ratio of 1.38 was observed (Frey et al., 2014a), and where *S. gotlandica*, a key player in denitrification coupled to S-oxidation, uses Nap for nitrate reduction, producing a denitrification Δδ^18^O:Δδ^15^N baseline of 0.5 (Frey et al., 2014b).

## 5. Summary and Conclusions

We investigated the drivers of dual nitrate N and O isotope effects associated to microbial nitrate reduction using a multifaceted approach involving experiments with pure cultures, natural microbial consortia of lake water, and in situ nitrate isotope analyses. The nitrate isotope data suggest that nitrification in the studied water column takes place under conditions where O atom exchange between nitrite and water is limited due to rapid nitrite turnover. In light of the commonly held perspective that nitrite oxygen isotopes in marine and freshwater denitrifying systems are completely equilibrated with the water O pool this begs further inquiries into the nitrate isotope systematics during nitrite oxidation/nitrate regeneration.

The biochemical and biogeochemical controls on the nitrate isotope signature in the natural environment are complex and are particularly complicated if Nar and Nap-driven nitrate reduction, nitrification, and anammox (not considered in this study) co-occur, all leaving their potentially superimposed fingerprints. Thus, the dual nitrate isotope approach has limitations in that it can only partly disentangle the combined signatures of simultaneous N-transformation reactions on the one hand, and the involvement of different nitrate reducing enzymes on the other. To study N-cycling processes in natural environments, coupled nitrate N and O isotope data are, therefore, best combined with quantitative information on *narG* and *napA* genes/transcripts, and on N-transformation rates. Nonetheless, our study confirms the usefulness of dual nitrate isotopes as a tool to provide information on N-cycling processes in net-denitrifying natural environments, for instance, to detect temporal changes in the relative importance of N-cycling processes as organotrophic denitrification, lithotrophic denitrification and nitrification.

## Data availability

Water column chemistry and experimental data are available through the Open Science Framework (https://osf.io/gfx7w/). Pure culture experimental data and treated 16S rRNA gene sequences (ASV) are deposited at the Zenodo repository (https://doi.org/10.5281/zenodo.14885067). Raw sequence data are made available at NCBI under the BioProjectID PRJNA772618 with the accession numbers SAMN47302969 through SAMN47303125 and under the BioProjectID PRJNA1234124 with the accession numbers SAMN47290646 through SAMN47290648.

## Acknowledgements

We thank Stefano Beatrizotti, Marco Simona, Fabio Lepori, Adeline Cojean-Egger, Guangyi Su and Maciej Bartosiewicz for their help during the sampling campaigns on Lake Lugano. We thank Sonia Tarnawski and Jeff A. Cole for providing *Pseudomonas* sp. and *Escherichia coli* strains, respectively. We also would like to thank the late María R. Miracle, Eduardo Vicente, and Xavier Sòria-Perpinyà for their help in making sampling of Lake La Cruz possible. Finally, we thank Thomas Kuhn for his excellent work in keeping the mass spectrometers running. This study was funded by the Swiss National Science Foundation project 153055 granted to JZ and MFL.

## Author contribution statement

JZ and MFL conceived the research project. JT, JZ, and MFL conceptualized research and experimental design. Pure culture experiments were conducted by JV, LB, and OR. JT, JZ, MFL, and EG conducted sampling campaigns and contributed to the data acquisition of physicochemical parameters in the water column of Lake Lugano. The incubation experiments with lake water were conducted by JT and analyzed by JT, MFL, JZ, and CF. JT measured and analyzed *in situ* nitrate isotopes. JT carried out the statistical analysis of nitrate isotope effects. JT wrote the paper, with substantial input from MFL, JZ, CF, and SDW. JT, JZ, and MFL are accountable for the integrity of the data, analysis, and presentation of the findings.

