## Supplementary Information for "Nitrogen and oxygen isotope effects during enzymatic nitrate reduction *in vitro* and by natural lake water consortia"

*The following supplementary material accompanies the manuscript*

### Supplementary tables:

**Table S1.** Selection of nitrate N and O isotope effects in pure-culture or enzyme-assay experiments and environmental water samples (marine water, groundwater, spring water, river water, lake water). SE = standard error, SD = standard deviation

| **Species, enzyme assay or study site** | **Nitrate reductase gene** | **$^{15}\varepsilon$ (Rayleigh) (‰)** | **$^{18}\varepsilon$ (Rayleigh) (‰)** | **$\Delta\delta^{18}O:\Delta\delta^{15}N$ or $^{18}\varepsilon:^{15}\varepsilon$** | **Reference** |
|---|---|---|---|---|---|
| *Ochrobactrum* sp. | *narG* | 7 to 23 | 7 to 23 | 0.94 to 1.02 | Granger et al. (2008) |
| *Pseudomonas chlororaphis* | *narG* | 11 to 30 | 16 to 21 | 0.83 to 1.02 | Granger et al. (2008); Kritee et al. (2012) |
| *Pseudomonas stutzeri* | *narG* | 5 to 20 | 5 to 18 | 0.86 to 0.92 | Granger et al. (2008) |
| *Paracoccus denitrificans* | *narG* | 9 to 33 | 17 to 33 | 0.80 to 1.04 | Granger et al. (2008); Kritee et al. (2012); Treibergs and Granger (2017) |
| *Marinobacter* sp. | *narG* | 15 to 23 | | 0.84 to 1.01 | Kritee et al. (2012) |
| *Thauera aromatica* | *narG* | 18 to 22 | 16 to 20 | 0.84 to 0.91 | Wunderlich et al. (2012) |
| *Aromatoleum aromaticum* | *narG* | 17 to 24 | 16 to 24 | 0.93 to 1.01 | Wunderlich et al. (2012) |
| *Thiobacillus denitrificans* | *narG* | 12.5 | 8.8 | 0.96 | Hosono et al. (2015) |
| *Pseudomonas aeruginosa* | *narG* | 24 to 29 | 20 to 26 | 0.85 to 0.91 | Asamoto et al. (2021) |
| *Bacillus vireti* | *narG* | 11 to 21 | 6 to 13 | 0.64 ± 0.04 SE | Asamoto et al. (2021) |
| *Bacillus bataviensis* | *narG* | 12 to 15 | 5 to 10 | 0.61 ± 0.06 SE | Asamoto et al. (2021) |
| *Acidovorax* sp. | *narG* | 19 to 25 | 18 to 24 | 0.97 to 1.03 | Chen et al. (2023) |
| *Pseudomonas* sp. | *narG* | 13 to 22 | 12 to 21 | 0.94 to 0.95 | this study |
| *Escherichia coli* | *narG* | 15 to 17 | 15 to 16 | 0.98 | this study |
| *Rhodobacter sphaeroides* | *napA* | 13 to 23 | 8 to 14 | 0.55 to 0.68 | Granger et al. (2008); Treibergs and Granger (2017) |
| *Sulfurimonas gotlandica* | *napA* | 13 to 28 | 8 to 14 | 0.43 to 0.68 | Frey et al. (2014b) |
| *viologen-fueled assay* | *napA* | 37 to 40 | 19 to 20 | 0.5 | Treibergs and Granger (2017) |
| *Pseudomonas aeruginosa* | *napA* | 32 to 35 | 15 to 18 | 0.59 ± 0.00 SE | Asamoto et al. (2021) |
| *Desulfovibrio desulfuricans* | *napA* | 22 to 25 | 14 to 16 | 0.63 ± 0.06 SE | Asamoto et al. (2021) |
| *Shewanella loihica* | *napA* | 21 to 22 | 12 | 0.55 ± 0.01 SE | Asamoto et al. (2021) |
| *Escherichia coli* | *napA* | 8 to 10 | 6 | 0.65 to 0.74 | this study |
| *Pseudogulbenkiania* sp. | *narG+napA* | 24 to 25 | 12 to 17 | 0.50 to 0.73 | Chen et al. (2020) |
| *Pseudomonas aeruginosa* | *narG+napA* | 23 to 29 | 14 to 27 | 0.63 to 0.97 | Asamoto et al. (2021) |
| *Paracoccus denitrificans* | *narG+napA* | 17 | 15 to 16 | 0.92 ± 0.01 SE | Asamoto et al. (2021) |
| *Pseudomonas* sp. | *narG+napA* | 8 to 30 | 8 to 28 | 0.90 to 0.99 | this study |
| Northern South China Sea | | | | 1 to 1.43 | Chen et al. (2019); Lao et al. (2019) |
| Xiangshen Bay | | | | >1 | Yang et al. (2018) |
| Upwelling Area Peru | | 19.2 | | 1 | Grasse et al. (2016) |
| Eastern North Pacific | | | | ~1.25 | Sigman et al. (2005) |
| Santa Barbara Basin | | 5 | | 1 | Sigman et al. (2003) |
| Baltic Sea | | 4.7 | 7 | 1.38 | Frey et al. (2014a) |
| Arabian Sea | | 34.5 | 28.8 | 1.2 | Gaye et al. (2013) |
| Szczecin Lagoon Outflow | | 3.5 | 4.8 | 1.2 | Korth et al. (2013) |
| Groundwater | | 27.6 | 18.3 | 0.67 | Mengis et al. (1999) |
| Groundwater | | 13.6 | 9.8 | 0.77 | Fukada et al. (2003) |
| Groundwater | | 5 to 17 | | 0.76 | Wexler et al. (2014) |
| Groundwater | | | | 0.5 to 1 | Soldatova et al. (2017) |

**Table S1 (continued).** Selection of nitrate N and O isotope effects in pure-culture or enzyme-assay experiments and environmental water samples (marine water, groundwater, spring water, river water, lake water). SE = standard error, SD = standard deviation

| **Species, enzyme assay or study site** | **Nitrate reductase gene** | **$^{15}\varepsilon$ (Rayleigh) (‰)** | **$^{18}\varepsilon$ (Rayleigh) (‰)** | **$\Delta\delta^{18}O:\Delta\delta^{15}N$ or $^{18}\varepsilon:^{15}\varepsilon$** | **Reference** |
|---|---|---|---|---|---|
| Groundwater | | 10 | | 0.42 to 0.72 | Bourke et al. (2019) |
| Groundwater | | | | 1.03 | Kang and Xu (2016) |
| Springs | | | | 0.59 | Heffernan et al. (2012) |
| Mississippi River | | 15.9 | 8 | 0.5 | Panno et al. (2006) |
| Ichetacknee River | | 3.1 to 5.6 | 1.6 to 3.7 | 0.87 to 1.11 | Cohen et al. (2012) |
| Lake Kizaki | | 24.4 ± 7.2 SD | 15.7 ± 3.7 SD | 0.64 | Sasaki et al. (2011) |
| Lake Okeechobee | | | | 0.59 to 0.72 | Ma et al. (2020) |
| Lake Bromont | | 31.2 | 53.3 | 1.7 | Botrel et al. (2017) |
| Tillari reservoir | | 8.7 | 10.7 | 0.85 to 0.95 | Bardhan et al. (2017) |
| Lake Lugano South Basin | | 11.2 | 6.6 | 0.57 | Lehmann et al. (2003) |
| Lake Lugano North Basin | | 9.1 ± 0.6 SE | 8.0 ± 0.8 SE | 0.89 ± 0.05 SE | Wenk et al. (2014) |
| Lake Lugano North Basin | | 4.8 to 13.5 | 4.8 to 13.8 | 0.67 to 2.15 | this study |
| Lake Lugano North Basin, incubations | | 13 to 26 | 9 to 24 | 0.60 to 1.04 | this study |
| Lake La Cruz, incubations | | 19 to 30 | 18 to 26 | 0.82 to 0.93 | this study |

**Table S2.** N vs. O isotope fractionation characteristics based on the analysis of in-situ nitrate samples from the redox transition zone of Lake Lugano North Basin. Outliers, i.e., isotope values resulting in a strong deviation from linear Rayleigh-type (for $^{15}\varepsilon$ and $^{18}\varepsilon$) and $\Delta\delta^{18}O$:$\Delta\delta^{15}N$ relationships (see Figure S8), were removed before determining the isotope effects. Values are given with the standard error of means.

| **Sampling campaign** | $^{15}\varepsilon$ **(‰)** | $^{18}\varepsilon$ **(‰)** | $\Delta\delta^{18}O$:$\Delta\delta^{15}N$ **(‰)** |
|---|---|---|---|
| February 2015 | 9.6 ± 0.3 | 10.1 ± 0.3 | 1.05 ± 0.03 |
| April 2015 | 4.7 ± 0.6 | 10.6 ± 0.8 | 2.15 ± 0.45 |
| June 2015[a] | - | 10.8 ± 1.0 | - |
| August 2015 | 10.0 ± 0.9 | 10.4 ± 0.9 | 1.40 ± 0.17 |
| October 2015 | 10.5 ± 0.4 | 13.7 ± 0.9 | 1.31 ± 0.07 |
| December 2015 | 11.4 ± 0.9 | 12.3 ± 0.8 | 1.06 ± 0.15 |
| March 2016 | 6.5 ± 0.6 | 12.9 ± 0.7 | 1.92 ± 0.29 |
| September 2016 | 13.5 ± 0.1 | 9.7 ± 1.3 | 0.91 ± 0.08 |
| November 2016 | 7.7 ± 1.4 | 4.8 ± 0.2 | 0.67 ± 0.08 |
| February 2017 | 5.9 ± 1.1 | 10.7 ± 2.1 | 1.80 ± 0.02 |
| October 2017 | 9.1 ± 0.5 | 11.3 ± 0.5 | 1.25 ± 0.01 |
| April 2018 | 8.4 ± 0.5 | 11.7 ± 1.4 | 1.41 ± 0.10 |
| **Average** | **8.8 ± 0.8** | **10.8 ± 0.6** | **1.36 ± 0.14** |

[a] The $^{15}\varepsilon$ value and the $\Delta\delta^{18}O$:$\Delta\delta^{15}N$ ratio were not determined due to the absence of a linear relationship in the Rayleigh plot and in the $\delta^{18}O$:$\delta^{15}N$ plot, respectively.

**Supplementary figures:**

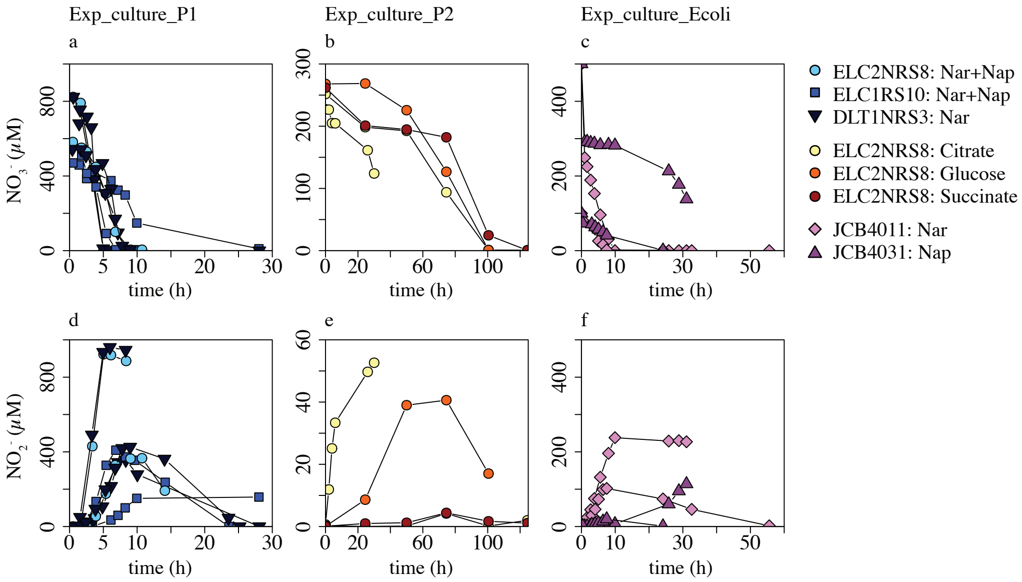


**Figure S1.** Concentrations of nitrate **(a-c)** and nitrite **(d-f)** in incubation experiments with *Pseudomonas* sp. and *Escherichia coli* cultures. **(a+d)** Exp_culture_P1 with *Pseudomonas* sp. strains DLT1NRS3, ELC2NRS8, and ELC1RS10, **(b+e)** Exp_culture_P2 with *Pseudomonas* sp. strain ELC2NRS8 growing with different carbon sources, and **(c+f)** Exp_culture_Ecoli with *E. coli* strains JCB4011 and JCB4031.

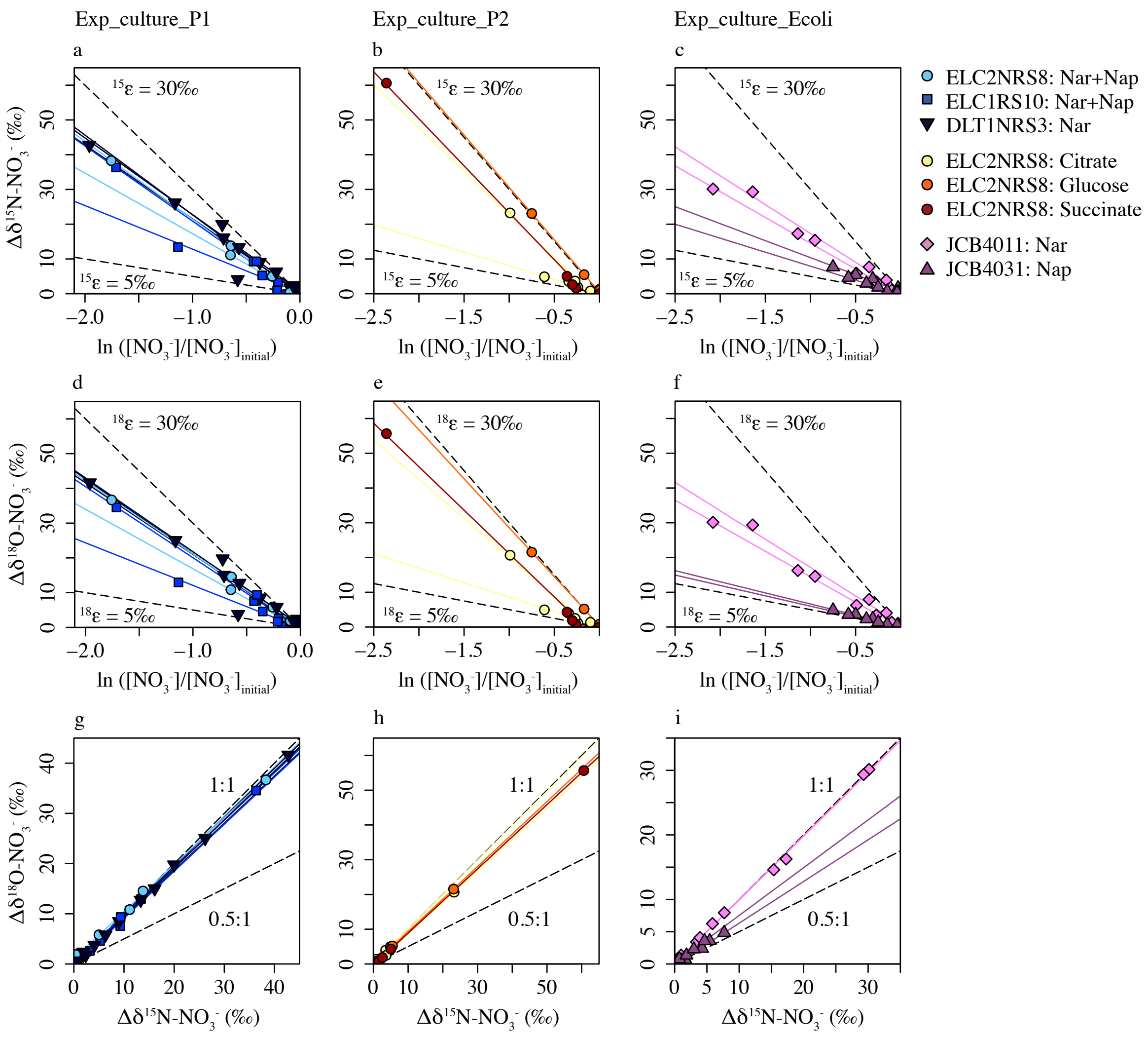


**Figure S2.** Isotope effects imparted by nitrate reduction in incubation experiments with *E. coli* and *Pseudomonas* sp. cultures, **a – c)** $\delta^{15}N$-Rayleigh plots, **d – f)** $\delta^{18}O$-Rayleigh plots, and **g – i)** $\Delta\delta^{18}O$-vs-$\Delta\delta^{15}N$ plots. **a), d),** and **g)** show results from Exp_culture_P1, **b), e),** and **h)** from Exp_culture_P2, and **c), f),** and **i)** from Exp_culture_Ecoli. Colored lines represent the linear regression of the respective experiment.

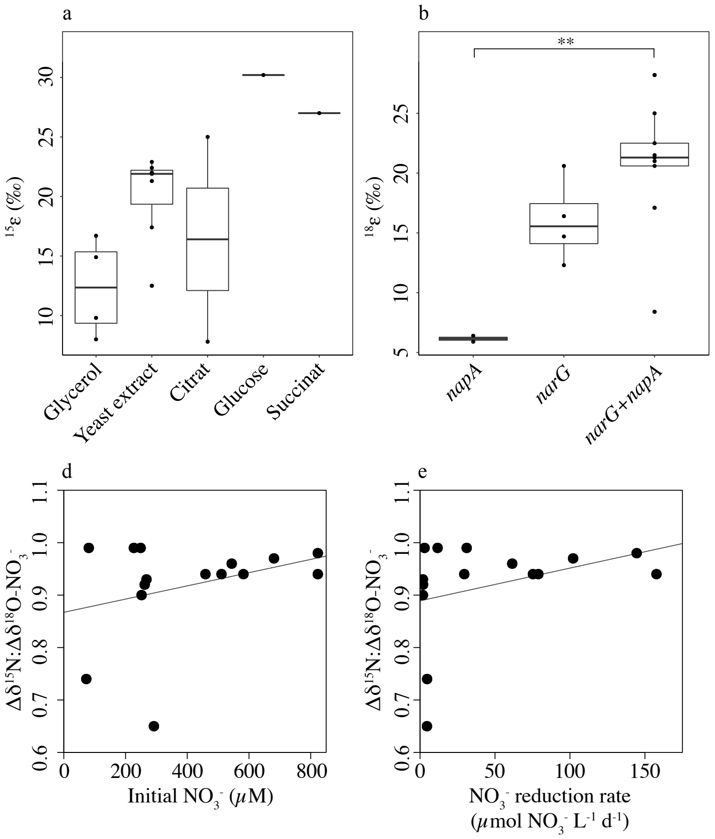


**Figure S3.** Variables and factors influencing $^{15}\varepsilon$, $^{18}\varepsilon$, and $\Delta\delta^{18}O:\Delta\delta^{15}N$ for nitrate reduction in pure-culture incubations (based on the final statistical models shown in Table 2). **a)** $^{15}\varepsilon$ depending on the carbon source, **b)** $^{18}\varepsilon$ depending on the nitrate reductase, **c)** $\Delta\delta^{18}O:\Delta\delta^{15}N$ depending on the type of nitrate reductase, **d)** $\Delta\delta^{18}O:\Delta\delta^{15}N$ depending on the initial $NO_3^-$ concentration, and **e)** $\Delta\delta^{18}O:\Delta\delta^{15}N$ depending on the $NO_3^-$ reduction rate. Stars indicate significance (*0.01 < p < 0.05, **0.001 < p < 0.01, ***$p$ < 0.001) of post-hoc Tukey tests.

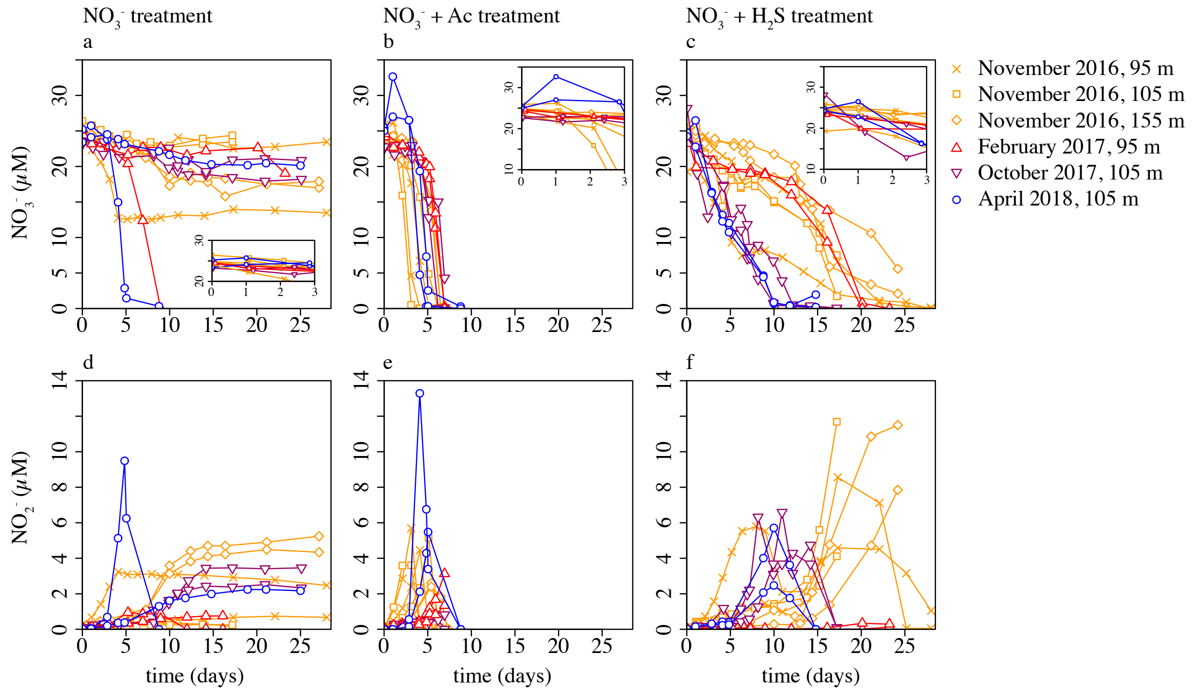


**Figure S4.** Concentrations of nitrate **(a-c)** and nitrite **(d-f)** in duplicate incubation experiments with natural net-denitrifying microbial consortia from Lake Lugano North Basin (Exp_LL) with added $NO_3^-$ **(a, d)**, $NO_3^-$ + acetate **(b, e)**, or $NO_3^-$ + $H_2S$ **(c, f)**. Inserts show nitrate concentrations within the first 3 days of the experiment. This graph has been previously published in Tischer et al. (2025). For nitrate reduction rates and isotope analysis, only time points with available isotope measurements were used.

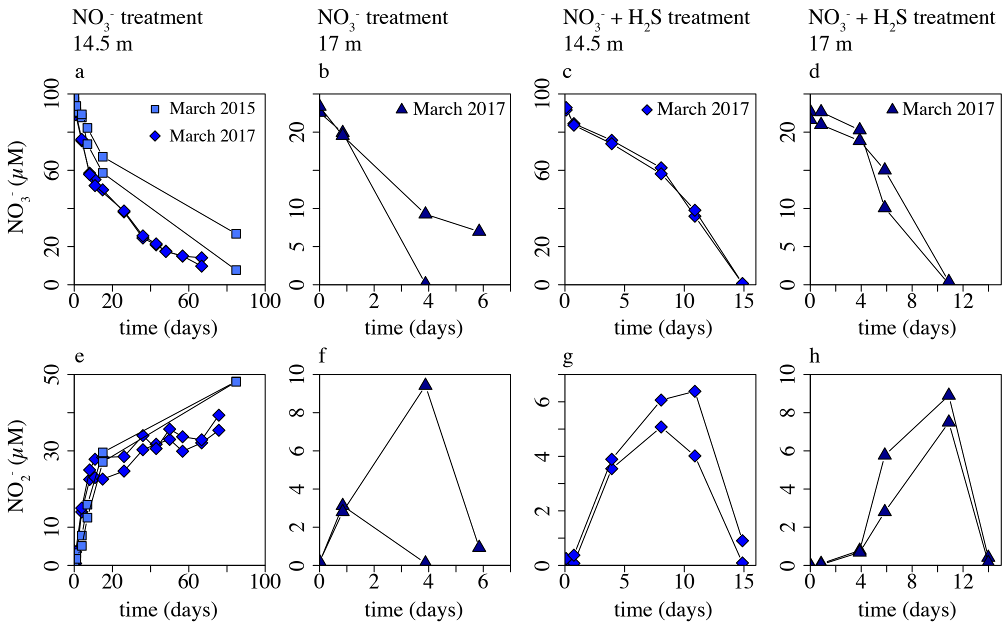


**Figure S5.** Concentrations of nitrate **(a-d)** and nitrite **(e-h)** in duplicate incubation experiments with natural net-denitrifying microbial consortia from Lake La Cruz (Exp_LC) with added $NO_3^-$ or with added $NO_3^-$ + $H_2S$.

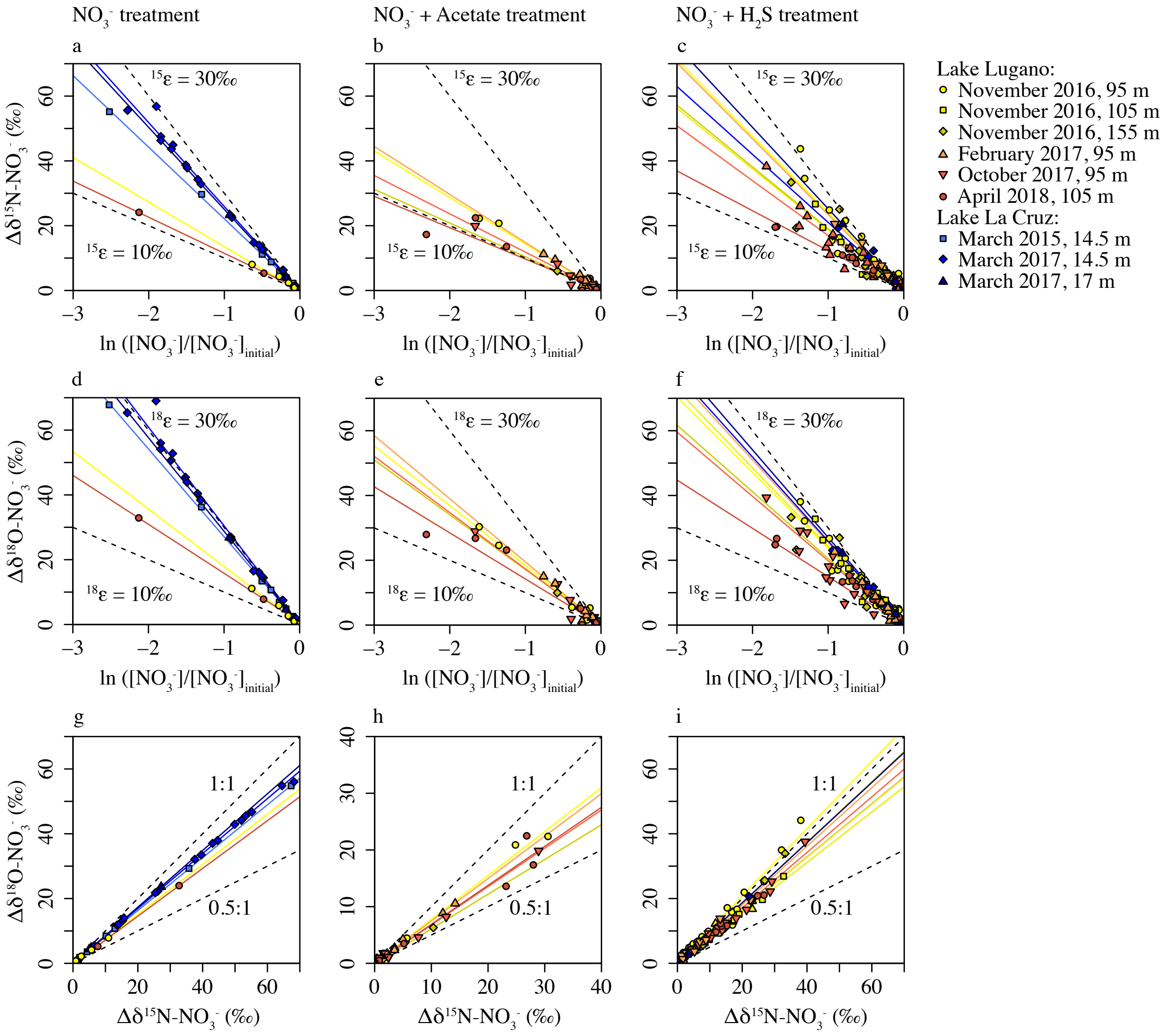


**Figure S6.** Nitrate isotope fractionation in incubation experiments with natural net-denitrifying microbial consortia from Lake Lugano and Lake La Cruz. **a – c)** $\delta^{15}$N-Rayleigh plots, **d – f)** $\delta^{18}$O-Rayleigh plots, and **g – i)** $\delta^{18}$O-vs-$\delta^{15}$N plots. **a), d),** and **g)** show results from the $NO_3^-$ treatment, **b), e),** and **h)** from the $NO_3^-$ + acetate treatment, and **c), f),** and **i)** from the $NO_3^-$ + $H_2S$ treatment. Colored lines represent the linear regression of the respective experiment.

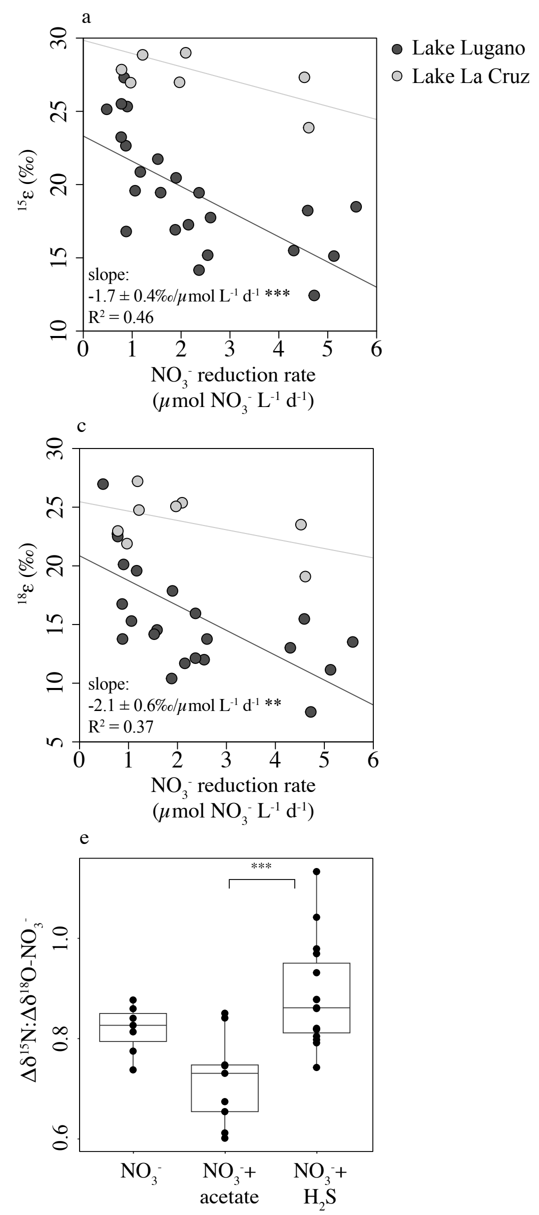


**Figure S7.** Variables and factors influencing $^{15}\varepsilon$, $^{18}\varepsilon$, and $\Delta\delta^{18}O:\Delta\delta^{15}N$ for nitrate reduction in incubation experiments with natural microbial consortia from the Lake Lugano North Basin and Lake La Cruz (statistical models shown in Table 4). **a)** $^{15}\varepsilon$ and **c)** $^{18}\varepsilon$ depending on the nitrate reduction rate and lake including the results of a linear regression model considering only the Lake Lugano incubations, **b)** $^{15}\varepsilon$ and **d)** $^{18}\varepsilon$ depending on the lake, and **e)** $\Delta\delta^{18}O:\Delta\delta^{15}N$ depending on substrates added. Stars indicate significance (*0.01 < p < 0.05, **0.001 < p < 0.01, ***$p$ < 0.001) of a linear regression model **(a, c)** or post-hoc Tukey tests **(b, d, e)**. Slopes (Lake Lugano only) are given with standard error of means and R squared ($R^2$).

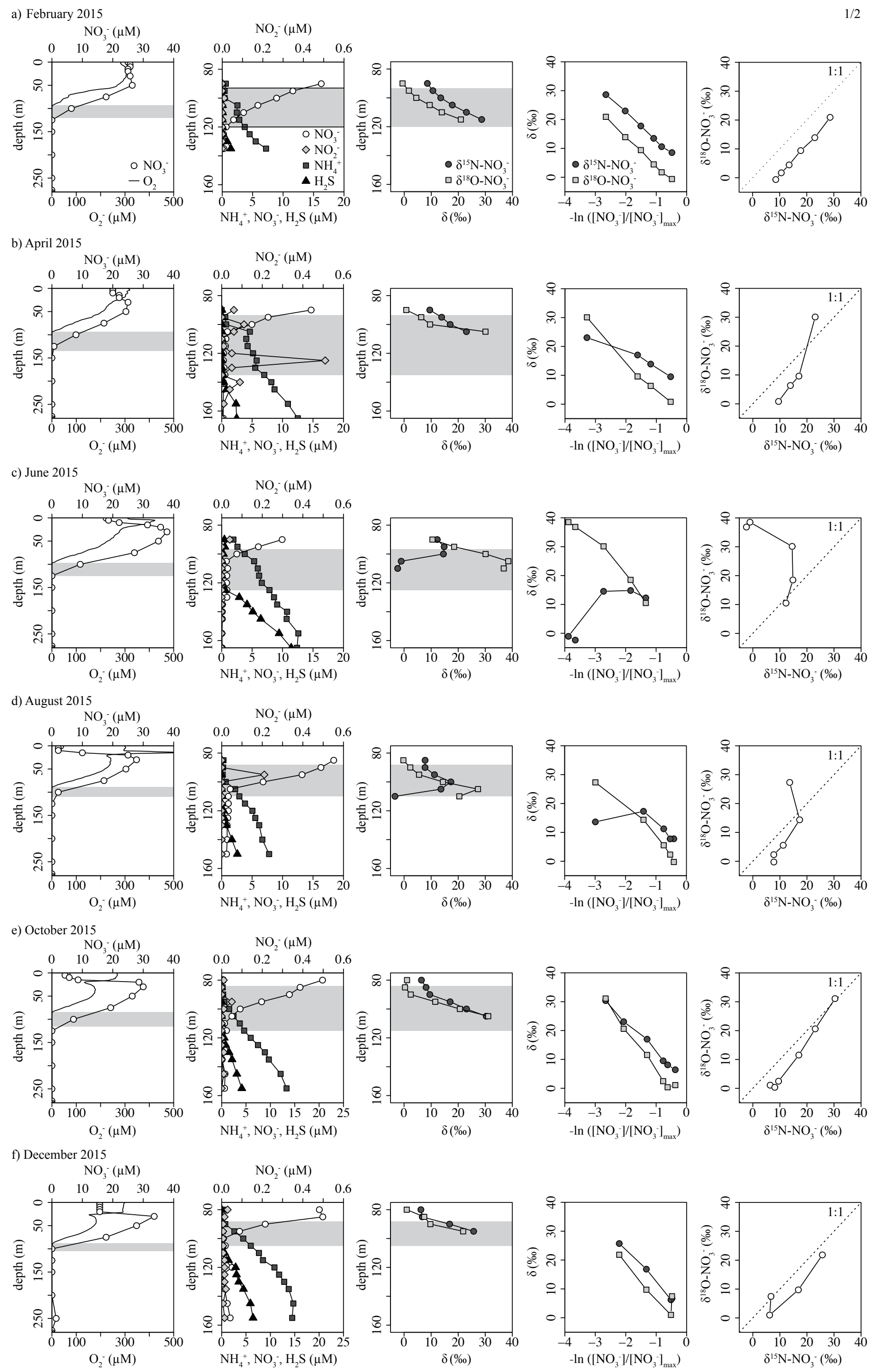


**Figure S8.** Water column profiles and nitrate concentration and isotope dynamics in Lake Lugano North Basin between February 2015 and April 2018. Columns from left to right: whole water column profiles of $O_2$ and $NO_3^-$ concentrations, $NO_3^-$ concentrations between 75 and 165 m depth of, profiles of $\delta^{15}N\text{-}NO_3^-$ and $\delta^{18}O\text{-}NO_3^-$, Rayleigh plots for $\delta^{15}N\text{-}NO_3^-$ and $\delta^{18}O\text{-}NO_3^-$, and $NO_3^-$ $\delta^{18}O$-vs-$\delta^{15}N$ plots. The redox transition zone is indicated by grey bars. Concentration data, except for February, June, and December 2015, have been previously published in Tischer et al. (2025).

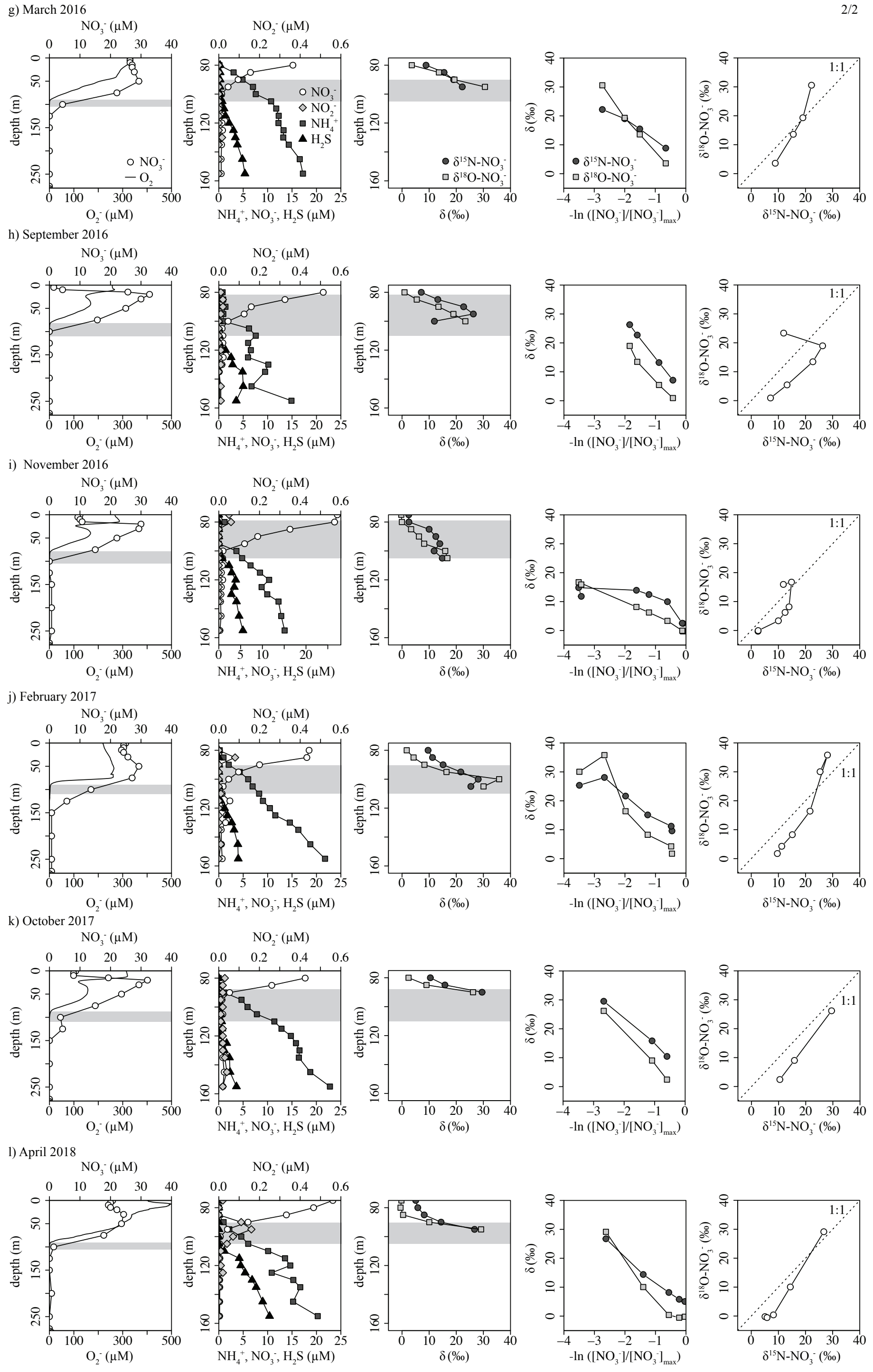


**Figure S8 (continued).** Water column profiles and nitrate concentration and isotope dynamics in Lake Lugano North Basin between February 2015 and April 2018. Columns from left to right: whole water column profiles of $O_2$ and $NO_3^-$ concentrations, $NO_3^-$ concentrations between 75 and 165 m depth of, profiles of $\delta^{15}N$-$NO_3^-$ and $\delta^{18}O$-$NO_3^-$, Rayleigh plots for $\delta^{15}N$-$NO_3^-$ and $\delta^{18}O$-$NO_3^-$, and $NO_3^-$ $\delta^{18}O$-vs-$\delta^{15}N$ plots. The redox transition zone is indicated by grey bars. Concentration data, except for February, June, and December 2015, have been previously published in Tischer et al. (2025).
